# Rapid acquisition of internal models of stimulus patterns by Purkinje cells enables faster behavioral responses

**DOI:** 10.1101/2021.04.28.441782

**Authors:** Sriram Narayanan, Aalok Varma, Vatsala Thirumalai

## Abstract

The cerebellum is known to optimize behavioral responses via error minimization, yet how past sensory experience shapes cerebellar output is still poorly understood. We imaged cerebellar Purkinje cells (PCs) in larval zebrafish during a behavioral assay in which they learnt to expect the direction of optic flow stimulus. PCs encoded signals of expectation and a graded error signal when a deviant stimulus was encountered. The amplitude of the error signal scaled based on stimulus history over minute-timescales. Granule cells (GC) did not encode error, however, some GC types encoded putative expectation signals. When PC population-wide readouts of expectation and error signals were strong, swim latency was low and vice versa, indicating participation of these signals in motor planning. Consistent with this, lesioning the cerebellum abolished expectation-dependent modulation of swim latency. Based on these results we propose that PCs use expectation and error signals to acquire and update internal models of the world.

**Highlights:**

- Larval zebrafish learn to expect repeatedly encountered optic flow stimuli
- Stimulus history over minutes contributes to an internal model of the sensory world
- Purkinje neuron activity encodes deviations from expected direction of optic flow
- The cerebellum uses learnt expectations to generate lower latency swim responses

## Introduction

A hallmark of animal behavior is the ability to learn fast and adapt to changing environments. With experience, animals across species can learn to expect events such as rewards or threats and predict the consequences of their actions ^1–3^. By integrating sensory evidence as and when available, the brain builds up an internal model of the world, which can be used to generate predictions to guide behavior. Neural mechanisms involved in this process often use error feedback to adaptively update models when the predictions they generate do not match outcomes in the real world ^4^. The cerebellum is a highly conserved brain region shown to be involved in this process but we do not yet have a circuit-wide understanding of the neural mechanisms involved. The primary input layer to the cerebellum is composed of a large number of granule cells (GCs), which are thought to expansively encode sensory and motor related inputs. Output from many GCs converge onto downstream Purkinje cells (PC). It is believed that by depressing GC synapses co-active with error signals from the inferior olive (IO), PCs learn to selectively respond to GC inputs that lead to error-free behavior ^5–7^.

Cerebellar function has been historically studied in the context of motor planning, learning and control ^8–13^. It is widely believed that with experience, the cerebellum learns how to control movements by acquiring predictive models of the motor system and the environment ^12,14,15^. Recently, the predictive functions of the cerebellum have been shown to be more generally applicable to non-motor domains, such as in encoding the expectation of rewards ^3,16–18^. Here, using a novel optomotor assay consisting of familiar and deviant optic flow stimuli, we probed whether the larval zebrafish cerebellum is involved in learning to expect stimuli based on prior experience over behaviorally relevant timescales of seconds to minutes.

The cerebellum is one of the oldest parts of the vertebrate brain, with cell types and connectivity well conserved in all vertebrates from fish to mammals ^19,20^. In larval zebrafish, the cerebellum is functional by 5 days post-fertilization (dpf) concomitant with the emergence of motor behaviors ^21^. At these stages, its small size and transparency allows the imaging of a substantial proportion of all cells in the cerebellum. By analysing the activity of both PCs and GCs in the same context from different fish, we hoped to arrive at a mechanistic description of cerebellar function. Previous studies have shown that the larval zebrafish cerebellum is involved in associative learning as seen in other species ^22^ and pre-motor activity in the fish cerebellum predicts future swim decisions ^2^, indicating a role in motor planning. PCs in larval zebrafish perform a combination of sensory and motor functions and are strongly activated by optic flow stimuli ^23,24^. The behavioral relevance of optic flow to fish means that no prior training is required, giving us the opportunity to probe innate mechanisms that enable them to learn what to expect from the sensory world.

Larval zebrafish sense optic flow and generate compensatory swims in the same direction (optomotor response or OMR) ^25–27^. The OMR is believed to help them stabilize their position in flowing water and can be reliably evoked in head-restrained fish by providing optic flow stimuli in the caudo-rostral direction on a screen placed beneath them ^28^. The latency with which optomotor swims are initiated depends strongly on the optic flow velocity ^29,30^. Apart from velocity, previous studies in both restrained and free swimming fish have also shown that the decision to swim is driven by a leaky integration of visual input, which is used to determine the direction of optic flow, especially in case of sparse and noisy environments ^31^. So far, these studies have looked at the OMR from a sensori-motor point of view and the contribution of learnt biases to this behavior has not been investigated. Here, we sought to ask whether the OMR is also influenced by a learnt expectation based on prior experience with optic flow stimuli.

In this study, we conducted a series of experiments on head-restrained larval zebrafish in a closed-loop optomotor environment, while simultaneously imaging calcium activity of specific cell types in the cerebellum. By introducing deviant stimuli after acclimatization to repetitive forward optic flow pulses, we probed whether fish learnt to expect the direction of optic flow stimulus. Our results indicate a causal role for the cerebellar circuit and directionally-tuned PCs in particular in signaling deviation from an expected pattern and in planning the motor response. We find that through this mechanism, larval zebrafish acquire internal models of their sensory world from experience, update these models when deviations occur and adjust their motor behavior accordingly.

## Results

### Purkinje cells (PCs) exhibit direction tuned responses to optic flow

To characterize the response properties of PCs to optic flow stimuli, we recorded calcium activity in head-restrained larval zebrafish swimming in a one dimensional closed-loop optomotor environment using two-photon microscopy (Figure 1A,B). Fish were initially presented with randomly interspersed pulses of 2s duration, of forward (+ve) or backward (-ve) moving gratings at 1cm/s or a ‘probe’ stimulus, where the gratings moved 1 pixel on the screen in the forward (+ve probe) or backward (-ve probe) direction, 4 times over the 2s (Figure 1C). Each stimulus was presented five times per fish with an interval of 5s between them (Figure 1C). We then fit the calcium responses of individual cells using forward, backward and motor regressors using multiple linear regression and selected cells whose responses could be fit with an `R`^2^ ≥ 0.6 (Figure 1D) ^23,32^. When we grouped cells based on the largest regression coefficient, we found that they segregated into either ‘forward’ or ‘backward’ tuned (Figure S1A) and there were no cells that were strongly activated by both directions of optic flow (Figure S1A,D). These results are consistent with previous findings which show that PC response to optic flow is direction specific ^23^. Fewer cells were activated the strongest during motor activity and not by optic flow stimuli. The motor correlated cells were also typically activated by forward optic flow stimuli (Figure 1E). This could be because of two reasons: since forward optic flow stimuli reliably evoked swims, PCs involved in generating the optomotor response could be activated by both sensory input as well as motor activity. Or, the strong correlation between motor activity and the forward optic flow stimulus could make it difficult to distinguish between them using regression.

**Figure 1:**
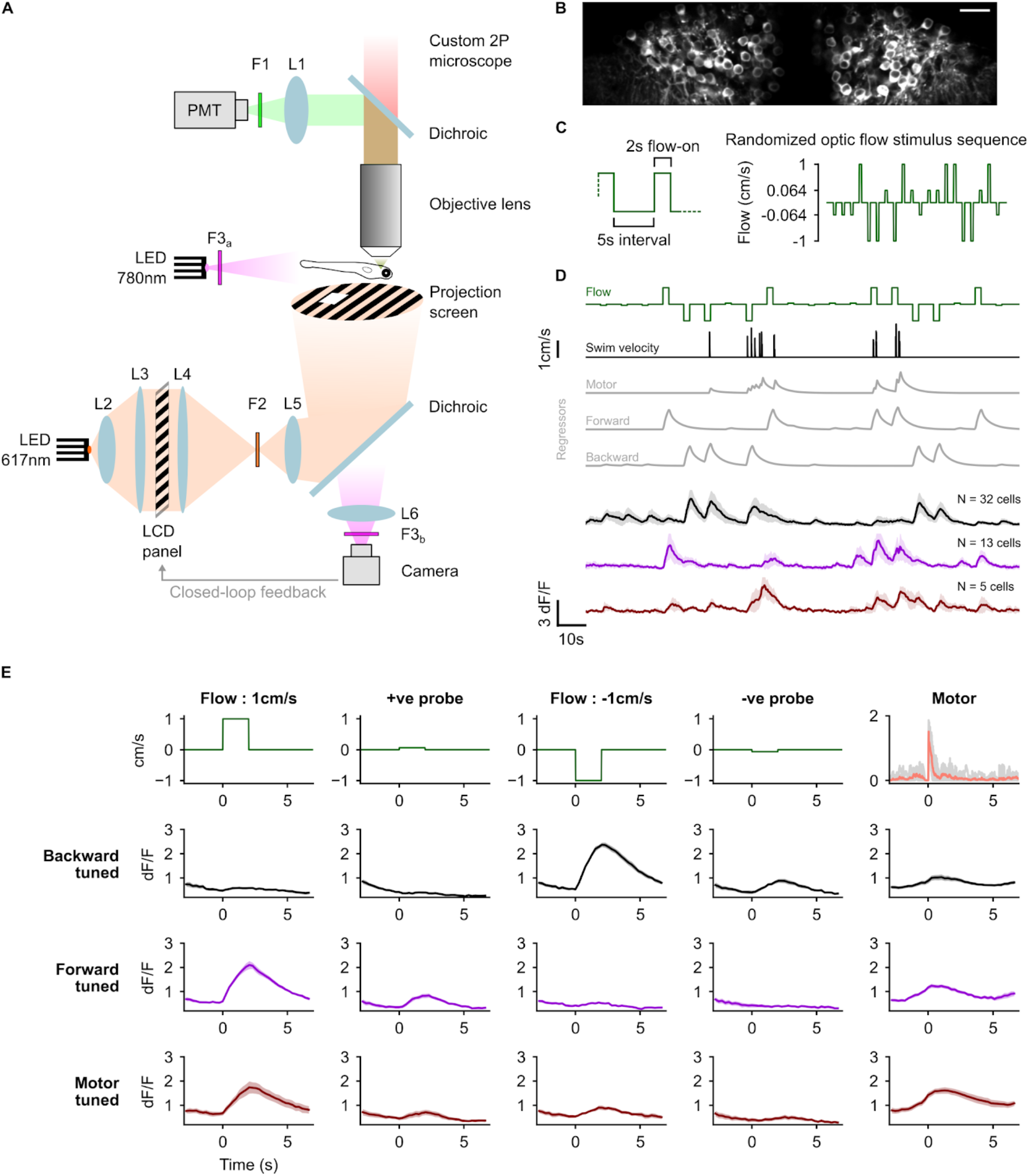
Purkinje cells show direction selective responses to optic flow stimuli. (A) Schematic of the experimental setup. (B) Average two-photon image of PCs expressing GCaMP6s from a representative fish. Scale bar : 20μm (C) Left: zoomed in view of the stimulus protocol highlighting the 5s interval between two stimulus pulses and the 2s pulse duration. Right: an example of the randomly ordered optic flow stimulus presented in the first trial. Note: the y-axis is scaled non-linearly in this schematic to highlight the small amplitude ‘probe stimulus’ pulses. (D) Classification of PCs based on tuning as backward (black), forward (magenta) or motor (brown) using regressors (gray) in a representative fish, shown as mean ± SD. (E) Average response profiles across fish of backward, forward and motor tuned cells (rows 2, 3 and 4 respectively) aligned to the start of each stimulus (column 1-4) or motor activity (column 5); shaded regions around the average calcium traces are SEM over N=8 fish. The first row shows the respective optic flow stimulus (columns 1-4) and average motor activity (column 5). Motor activity traces in gray represent the average for individual fish.

We then proceeded to classify cells as either ‘forward’, ‘backward’ or ‘motor’ based on the largest regression coefficient and strong correlation (Pearson’s r ≥ 0.6) with the respective regressor. Using this approach, we were able to classify 24±12% (mean ± SD) of all cells in one of the three groups. This classification was used in subsequent trials to group cells based on their tuning. Optic flow stimuli in both the forward (+ve) and backward (-ve) directions at 1cm/s caused strong activation in distinct populations of PCs (Figure 1E column 1,3). Both the positive and negative probe stimuli, being on average 15 times weaker than the 1cm/s optic flow stimuli, caused a much lower but detectable activation in PCs (Figure 1E column 2,4). Responses to the probe stimuli were also direction tuned, even though these stimuli consisted only of suggestive 1 pixel shifts in grating position, corresponding to a 3.2° visual angle, in the forward or backward direction (Figure 1E column 2,4). Forward moving gratings at 1cm/s reliably evoked swims. There were rarely any swims in response to the probe stimuli. Backward moving gratings at 1cm/s either failed to induce swim responses or caused struggles and/or large amplitude turns and hence were not presented subsequently.

### Response to optic flow in PCs is modulated by stimulus history

After the first trial where stimuli were presented in random order (Figures 1C, 2A), subsequent trials in the next phase of the experiment were conducted to test whether stimulus history influences PC response to deviant optic flow stimuli. We acclimatized fish with 8 repetitions of 2s long forward optic flow stimuli at 1cm/s, which consistently evoked slow forward swims in closed-loop (Figure 2B-E). The trials then proceeded into a probe phase where different probe stimuli were presented as shown in Figure 2A. These stimuli were either 3 pulses of the negative probe stimulus, 3 pulses of the positive probe stimulus or stationary gratings (no probe). In addition to these trials, a fourth trial was included where acclimatization pulses were presented at randomized time intervals (range: 1.5-8.5s, mean: 5s) instead of the fixed 5s interval. This was followed by 3 pulses of the negative probe stimulus. Each of the trials were presented three times in a different randomized order for every fish and the responses were averaged by trial type for each cell. We then analyzed the effect of acclimatization on the response to probe stimuli, with respect to baseline responses measured when stimuli were presented in random order in the first trial.

**Figure 2:**
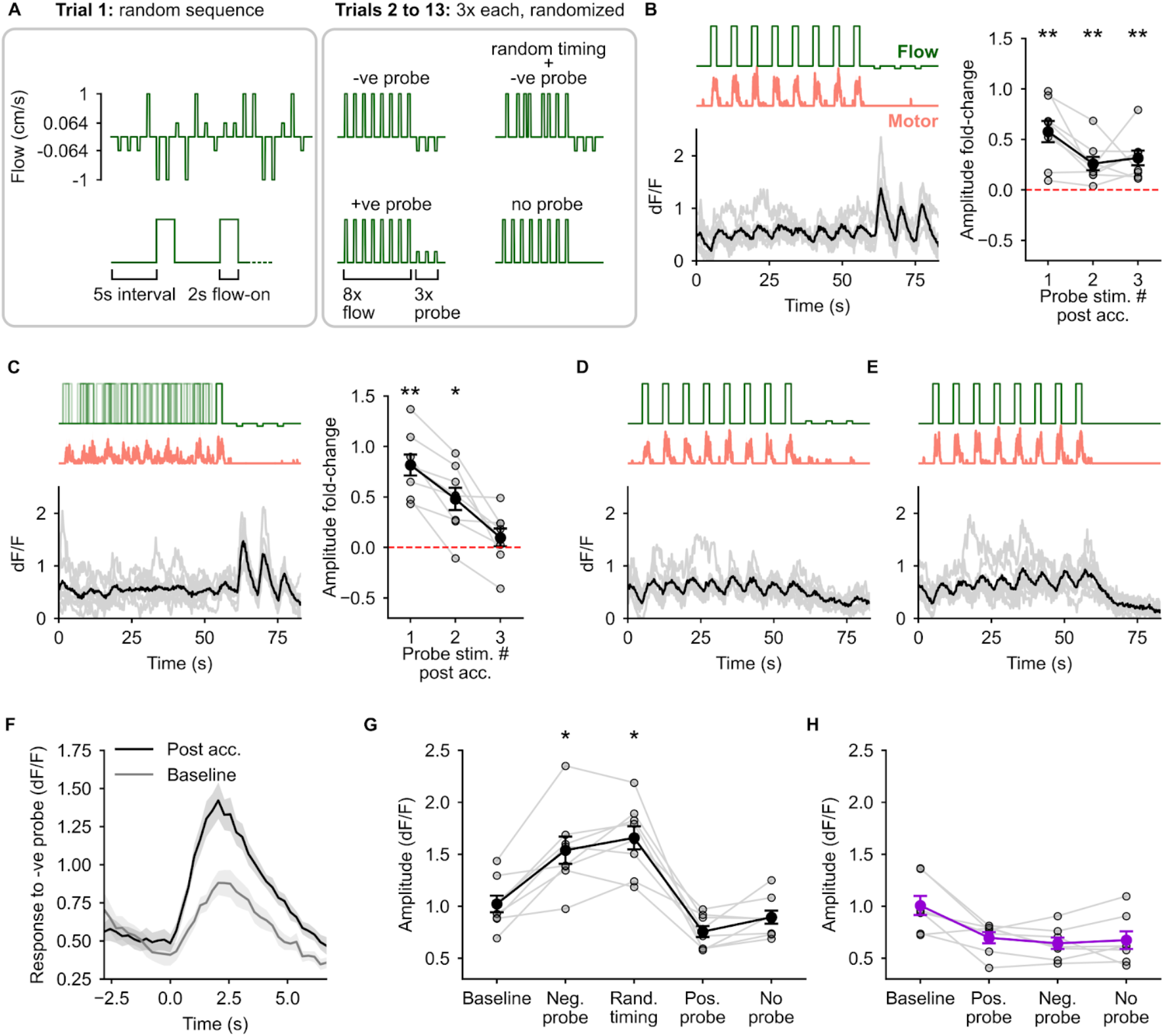
A change in optic flow direction is encoded in enhanced responses to probe stimuli. (A) Schematic of the experimental protocol. Trials 2-13 began with an acclimatization phase of 8x 1cm/s optic flow pulses followed by a probe phase. (B-E) Average calcium responses in the backward tuned cells to the different trials (B: negative probe, C: random stimulus timing followed by negative probe, D: positive probe and E: no probe). The respective optic flow stimuli and average motor activity are shown above. Individual calcium traces in gray are the responses of individual fish (N=8). Fold-change in peak amplitude in response to the three presentations of the negative probe stimuli w.r.t the baseline amplitude in trial 1 are shown for (B) and (C). ** : p < 0.01 and * : p < 0.05 by Wilcoxon signed-rank test showing an elevation in amplitude w.r.t baseline (dashed line at 0). There were no large calcium responses in the backward tuned cells to the positive probe or the zero probe (D, E). (F) Average calcium response in backward tuned cells to the negative probe stimulus when presented in random order in trial 1 (gray) and to the first presentation of the same stimulus post acclimatization (black) showing the enhancement of the calcium response post acclimatization. (G, H) The peak amplitude in dF/F of the trial averaged calcium response to the respective stimuli shown on the x-axis for the backward tuned cells (G) and the forward tuned cells (H). * : p < 0.05 w.r.t the baseline amplitude by post-hoc Conover’s test following a significant Friedman’s test (p = 3×10^-5^). N=8 fish in (G), N=7 fish in (H). No large calcium responses were seen in forward tuned cells to any of the probe stimuli (Figure S2) and hence no statistical analysis was performed on the amplitude values.

In backward-tuned cells, post acclimatization, the response to negative probe stimulus showed a large amplitude increase compared to baseline (Figure 2B,F,G). This enhanced response exhibited a decay over multiple presentations of the probe stimulus and approached baseline levels (Figure 2B,C). The increase in response amplitude was present to an equivalent extent when the timing of the acclimatization stimulus was randomized, indicating that timing did not play a role in the enhancement of the probe response (Figure 2C,G). Positive probe stimuli evoked no detectable calcium events in these cells (Figure 2D,G). In forward-tuned cells, negative probe stimuli evoked no response, whereas, positive probe stimuli showed no enhancement with respect to baseline (Figure 2H, S2). Both forward tuned and backward tuned cells did not show any responses when stationary gratings were presented in the probe phase (Figures 2E,G,H, S2C). Collectively, these results point to history-dependent enhancement of directionally-tuned PC responses. Specifically, in backward-tuned PCs, weak backward flows during random presentation evoked a small response; the same stimulus after the fish was acclimatized with forward flows evoked an enhanced response in the same PCs. It is possible that the enhanced response to the probe stimulus encodes a deviation from expected optic flow direction.

### Self generated change in optic flow due to closed-loop motor activity does not cause strong PC activation

Motor activity in closed-loop often caused a sharp change in optic flow direction. We asked if the enhanced probe response was also observed when the fish’s own swimming resulted in comparable or larger backward grating movement. If a self-induced reversal of optic flow direction caused a comparable activation of PCs as the negative probe stimulus, it would indicate that the response simply encoded a sensory change irrespective of what caused it. On the other hand, if the enhanced PC response to the negative probe stimulus indeed represented an unexpected externally imposed change, we would not expect to see strong activation in backward tuned PCs.

Forward moving gratings at 1cm/s reliably evoked swims. Occasional swim bouts were observed in response to the positive probe stimulus in some fish. Both the negative probe stimulus and stationary grating did not evoke swim responses (Figure 3A,B). In closed-loop, the net optic flow stimulus presented to the fish is the difference between the externally provided optic flow stimulus and the virtual swim velocity of the fish (Figure 3C) (See Methods). On average, the net optic flow stimulus seen by the fish showed a sharp dip below zero during swim bouts (Figure 3D) and the amplitude of the negative component was much greater during swims than the amplitude of the probe stimulus (Figure 3D,E). The amplitude of the calcium response in backward tuned cells to the motor activity triggered reversal of optic flow direction was negligible compared to the response to the negative probe stimulus (Figure 3D,E).

**Figure 3:**
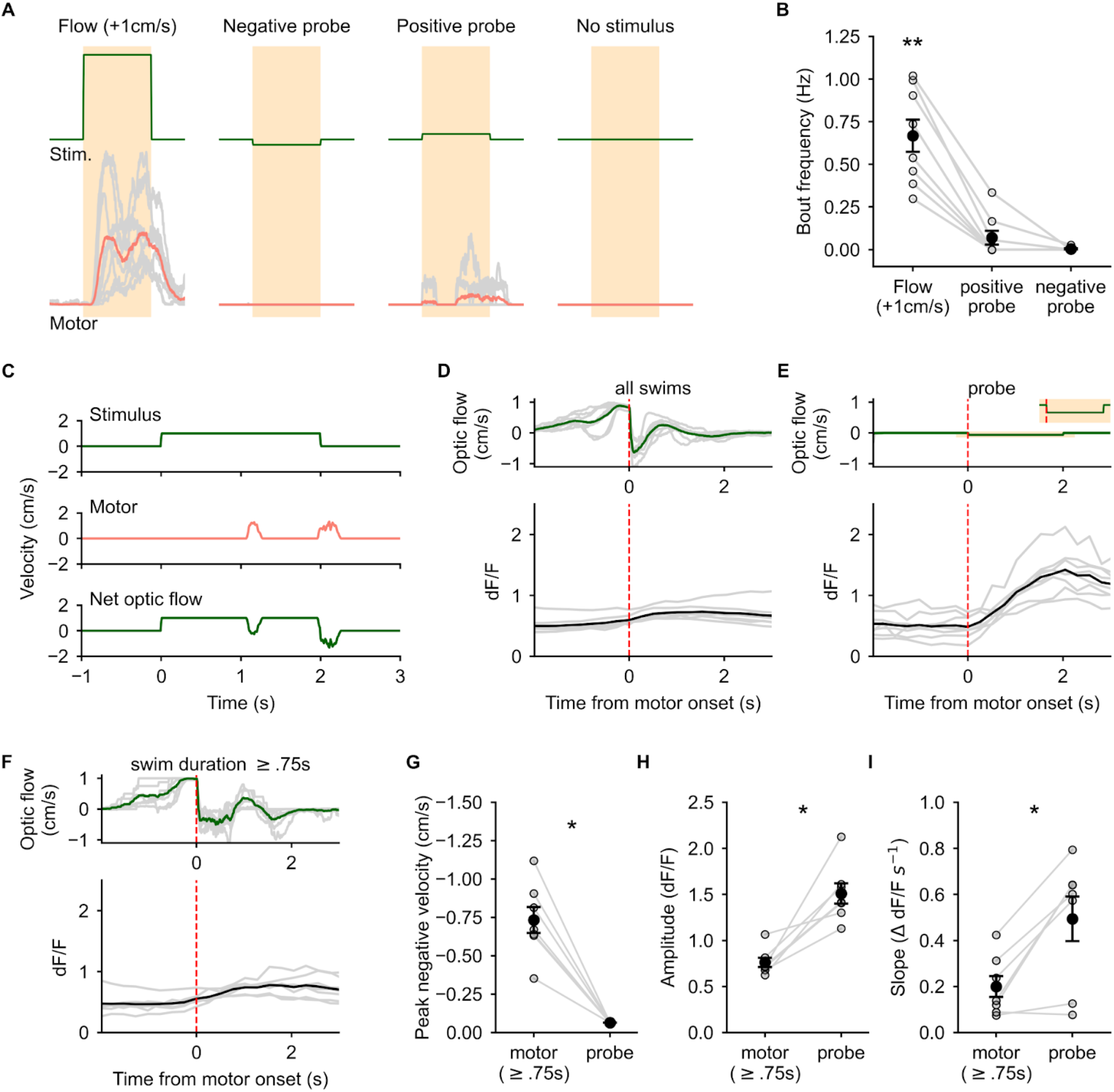
Self induced change in optic flow direction does not cause PC activation. (A) Average motor activity (bottom, red) during 1cm/s forward optic flow, negative probe, positive probe and no optic flow stimuli (top, green). (B) Forward optic flow reliably evoked swims, whereas very few and no swims were seen in response to positive probe and negative probe stimuli respectively. N=8 fish, ** : p = 0.0006 by Friedman’s test, p = 0.011 (forward flow vs +ve probe) and p = 0.002 (forward flow vs -ve probe) by post-hoc Conover’s test. (C) Example 1cm/s forward optic flow stimulus (top), motor activity (middle) and the resultant net optic flow experienced by the fish in closed-loop (bottom). (D) Average optic flow experienced by the fish during all swim bouts (top) and co-aligned calcium responses in backward tuned PCs (bottom). (E) Average optic flow stimulus experienced by the fish during negative probe pulses (zoomed inset shows the probe pulse) (top) and co-aligned calcium responses in backward tuned PCs (bottom). Gray traces represent the average response for individual fish. (F) Average optic flow experienced by the fish during swim bouts ≥ 0.75s in duration (top) and co-aligned calcium responses in backward tuned PCs (bottom). (G) Magnitude of the negative component of optic flow during swims is much larger than the amplitude of the negative probe stimulus. (H) Peak amplitude of the calcium response in backward tuned cells to the negative probe stimulus is much larger than the response to swim bouts. (I) The slope of the rise phase of the calcium response in backward tuned cells is significantly steeper for the negative probe stimulus than during motor activity. N=7 fish, * : p = 0.016 (G, H) and 0.03 (I) by Wilcoxon signed-rank test.

To compare the two, we selected swim bouts that lasted 750ms or more, corresponding to 3 calcium imaging frames, providing a sizable window for comparison of calcium activity. The average amplitude of the self-generated negative optic flow was much greater than the negative probe amplitude (Figure 3F,G). However, the amplitude of the calcium signal was greater in response to the probe stimulus than during swim events longer than 750ms (Figure 3H). Similarly, the slope of the calcium signal measured over 3 imaging frames from the onset of the probe stimulus was also greater than that in the case of swim events (Figure 3I).

These results show that sensory tuned PCs are not activated by self-generated sensory input. The enhanced response to the negative probe stimulus indeed encodes an externally triggered deviation in optic flow direction.

### PC response to an abrupt change in optic flow stimulus encodes deviation from expectation

If fish indeed learnt what stimulus to expect, we hypothesized that the level of confidence they have in what to expect from the sensory world would be reflected in a correspondingly scaled enhancement of the PC response to deviant stimuli. To test this hypothesis, we designed an experiment where we attempted to induce varying levels of confidence by varying the number of repetitions of the acclimatization stimulus. In this experiment, in trial 1, we presented forward and backward flows in random sequence as we did before (Figure 1C). Following this, we paired behaviorally salient 1cm/s forward optic flow with the negative probe stimulus. Fish were presented with a set of four trials, repeated 3x each, in random order as shown in Figure 4A. Each trial began with a pre-acclimatization phase (‘*pre-acc*’) of 3 pulses of the negative probe stimulus, followed by an acclimatization phase of either 2, 4, 8, or 16 pulses of forward moving gratings, followed by a post-acclimatization phase (‘*post-acc*’) of 3 pulses of the negative probe stimulus (Figure 4A). The trials were separated by a 30s inter-trial interval (ITI) during which stationary gratings were presented. We identified backward-tuned PCs from their responses during trial 1 and then analyzed the *pre-acc* and *post-acc* probe responses of these cells after 16, 8, 4 or 2 acclimatization pulses.

**Figure 4:**
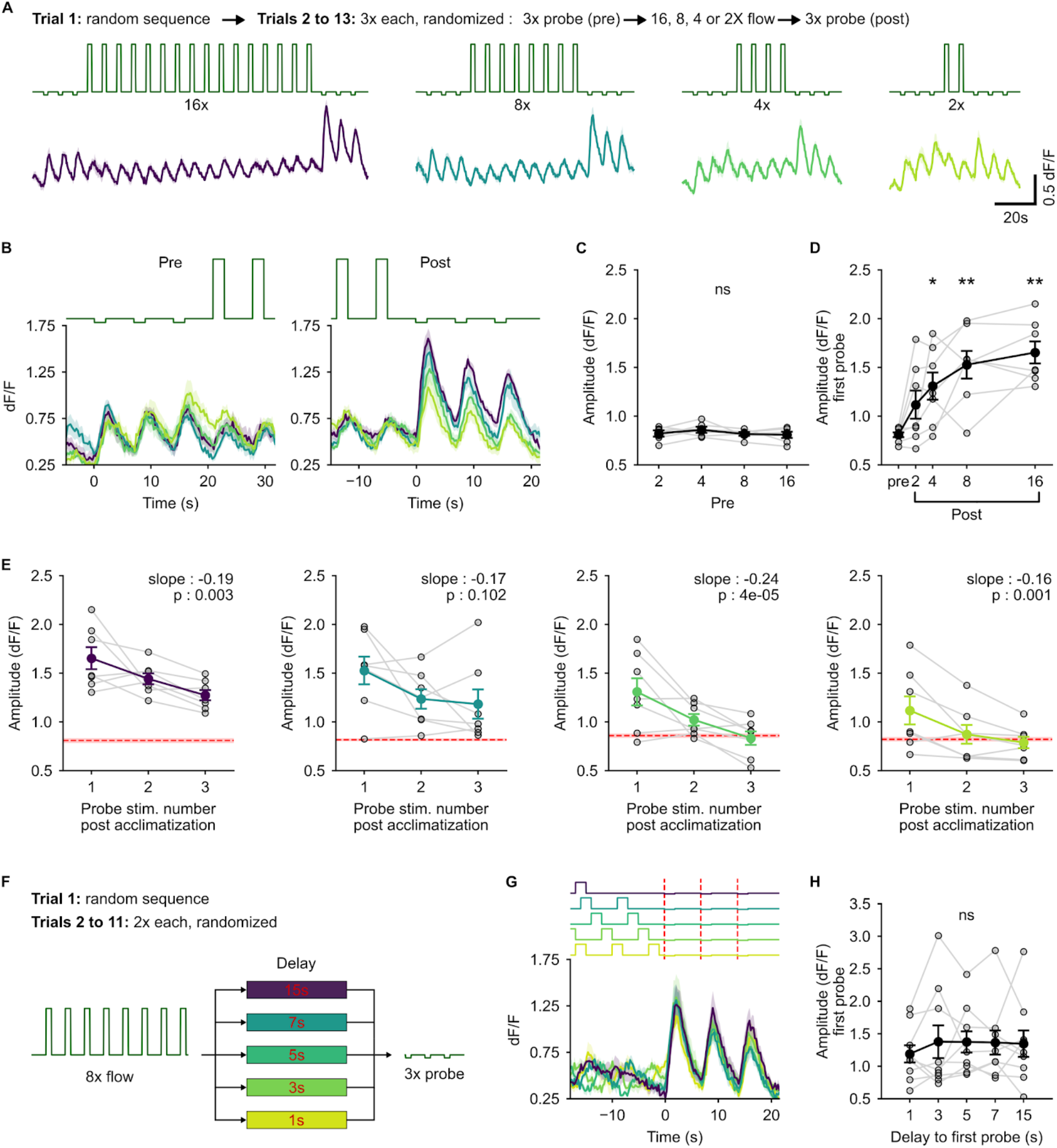
Probe response in PCs encodes deviation from the expected direction of optic flow. (A) Top (text): description of the experimental protocol. Middle: 3x probe pulses presented before (pre) and after (post) acclimatization pulses of forward optic flow at 1 cm/s. The number of acclimatization pulses was varied (16, 8, 4 or 2) and these blocks were presented in randomized order. Bottom: Corresponding calcium responses in backward tuned cells. Solid line is the mean for N=7 fish and shaded regions are SEM. Colors indicate the number of acclimatization pulses. (B) Zoomed in view of the average calcium response of backward tuned cells to pre- (left) and post-probe (right) pulses aligned to the onset of the first probe. (C) Average amplitude of the response to pre-probe stimuli. ns : p = 0.54. The 3 probe pulses were pooled as there was no difference between them. (D) Average amplitude of the response to the first post-probe pulse scales with the number of acclimatization pulses. p = 0.0015 by Friedman’s test, * : p < 0.05, ** : p < 0.01 by post-hoc Conover’s test w.r.t the pre-probe amplitude. (E) Decay of the amplitude of the enhanced response to the post-probe stimulus over the three successive post-probe pulses for the different number of acclimatization pulses. Red dashed line with the shaded region shows the mean and SEM of the respective pre-probe amplitudes. Slope and p-value by linear mixed-effects are inset in each plot. (F) Schematic of the experimental protocol for the delayed probe experiments. 3x probe pulses were presented with a variable delay from 1s to 15s after 8x acclimatization pulses. (G) Average response of the backward tuned cells aligned to the onset of the probe pulses. Shaded regions are SEM across N=9 fish. Colors indicate the delay as shown in (F). (H) Amplitude of the response to the first probe pulse does not depend on the delay. ns : p = 0.98 by Friedman’s test.

There was no change in response amplitude across all trials in the *pre-acc* phase (Figure 4C). The *post-acc* responses indeed scaled with the number of acclimatization pulses (Figure 4A-D). The response amplitude to the first probe pulse in the post phase increased on average 2-fold with increasing number of repetitions of the acclimatization stimulus. The scaling was non-linear and the population average followed a logarithmic trend. The increase in amplitude was visually apparent starting from 2 acclimatization pulses and became statistically significant with 4 or more pulses. The scaling approached saturation at 16 acclimatization pulses (Figure 4D). To rule out the possibility that the scaling depended only on the elapsed time between the *pre-acc* phase and the *post-acc* phase, we looked at the effect of the 35s interval (30s ITI + 5s inter-pulse delay) between the end of a trial and the first pre-probe pulse of the next trial and compared that to the scaling caused by the 4-pulse acclimatization, which spanned an equivalent duration (33s between probe pulses) (Figure S3A,C). Stationary gratings presented during the ITI did not lead to an enhancement of the response compared to the 4-pulse acclimatization (Figure S3B,D,E), nor was the response post stationary gratings significantly different from the baseline response to the negative probe stimulus when presented in random order (Figure S3F). These results indicate that the enhancement of PC response to probe depends on repetitive experience with the acclimatization stimulus, suggesting that fish have formed an experience based expectation of what stimulus they are likely to encounter.

A possible explanation is that the enhanced responses to deviant stimuli could be the result of accumulation of stimulus history by integration of sensory input with long time constants, representing a prior belief in what stimulus to expect. Future inputs are then evaluated with respect to this prior belief, leading to enhanced responses when deviations are encountered. To test this hypothesis, we presented a separate set of fish with a constant 1cm/s forward optic flow stimulus in the acclimatization phase, for an equivalent time duration as in the pulsed acclimatization experiments (Figure S4A). A constant optic flow acclimatization stimulus did not lead to an enhanced *post-acc* response with respect to *pre-acc* responses (Figure S4B-D) and the post responses were significantly lower in amplitude than the corresponding pulsed acclimatization stimuli (Figure S4E). These results show that the underlying mechanism is not based on a simple integration of sensory input.

Alternatively, the enhanced response to deviant stimuli could be based on an experience gated updating of what to expect from the sensory world. If this were indeed the case, we expected two properties to be exhibited by the *post-acc* response. First, multiple presentations of the probe stimulus would lead to a decay of the enhanced response till it comes back to the baseline sensory response amplitude. And second, the degree of enhancement of the response to the first *post-acc* stimulus must be independent of delay from the acclimatization phase as no updates to the expectation have been made during the delay period.

When we looked at the *post-acc* responses, there was indeed a significant decay in amplitude from the first to the third presentation of the probe stimulus. The amplitude of the *post-acc* response decayed back to the *pre-acc* levels in the 4-pulse and 2-pulse acclimatization trials (Figure 4E). Next, to test whether the enhancement of the response to probe decayed on its own or if it required fish to experience the probe stimulus, we conducted a separate experiment on a different group of fish. No pre-acclimatization probe pulses were presented in this experiment. Fish were acclimatized to 8 pulses of 1cm/s forward optic flow, followed by 3 probe stimulus pulses, separated by 5s each. The delay to the first probe pulse from the last acclimatization pulse was varied from 1 to 15s as shown in Figure 4F. The amplitude of the response to the first probe pulse was independent of the delay from the acclimatization phase (Figure 4G,H) and exhibited a decay over the three repeats of the probe stimulus (Figures 4G, S5). These results show that the expectation is updated with experience and does not decay with time, within the limits of the 15s window. There being no change in amplitude of the response to the first probe also suggests that fish learn what stimulus they are likely to encounter rather than the time at which it would occur.

Taken together, the above results suggest that larval zebrafish learn to expect optic flow stimuli they are likely to encounter based on stimulus history over minute-timescales. A deviant probe stimulus caused enhanced responses in PCs, which scaled with the number of repetitions of the acclimatization stimulus. This enhanced response decayed back to baseline with repeated presentations of the probe stimulus and represents an error in expectation, which is likely to play a role in updating an internal model of the sensory world as new evidence is encountered.

### Cerebellar granule cells (GCs) encode sensory input but not deviation from expectation

PCs receive excitatory inputs from two distinct synaptic pathways, namely parallel fibers (PFs) from cerebellar granule cells (GCs) and climbing fibers (CFs) from the inferior olive (IO). It is believed that GCs encode sensory context and motor activity whereas the CF pathway carries reinforcing error signals. PF synapses co-active with the error signal are suppressed and those de-correlated with error selectively contribute to PC activity, subsequently affecting behavior. According to this framework of cerebellar function, we would expect GCs to represent an array of sensory tuned responses but not the error in expected optic flow direction.

To identify the input pathway(s) to PCs encoding deviation from the expected direction of optic flow, we performed serial volumetric calcium imaging of the GC population expressing GCaMP6f under the *neurod* promoter (Figures 5A,B). A single hemisphere of the cerebellum was densely sampled across 6 imaging planes separated by 10μm, yielding a total of 6465 GCs across 8 fish. This accounts for ~27% of all GCs in one hemisphere of the 7-day old larval zebrafish cerebellum. The stimulus protocol consisted of two trials separated by 30s. The first trial was a randomized sequence of stimulus pulses to determine baseline responses of GCs to optic flow. The second trial began with 8 acclimatization pulses of 1cm/s followed by 3 negative probe pulses, then 3 pulses of 1cm/s and ended with an equivalent period of stationary grating (Figure 5A).

**Figure 5:**
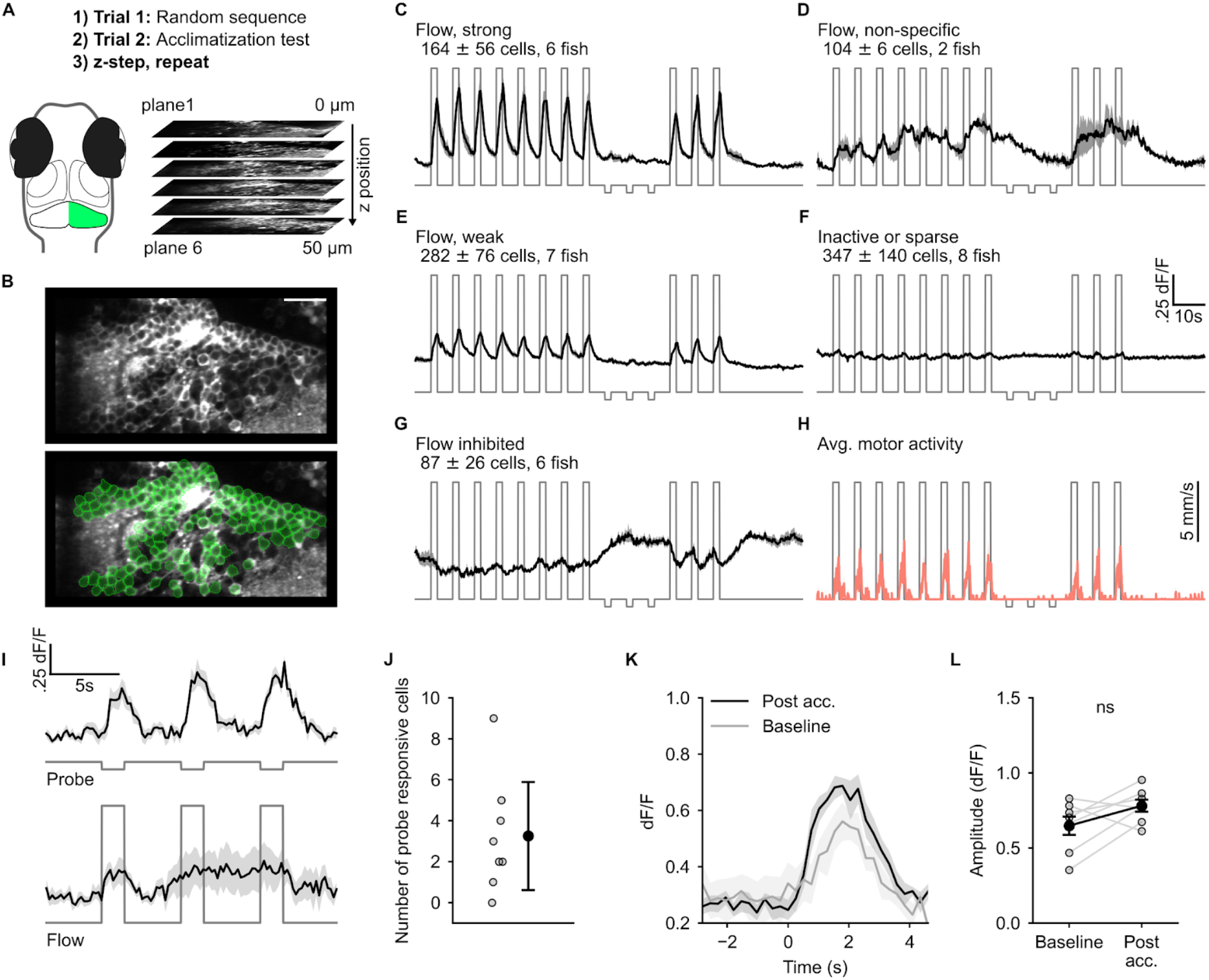
Granule cell responses reveal absence of the error signal. (A) Schematic of the experimental protocol. (B) A representative imaging plane (top) and ROIs drawn around GC cell bodies using semi-automated anatomical segmentation (bottom). (C-G) Functional response types identified using correlation based clustering. The cluster label, number of cells (mean ± SD) and the number of fish in which the cluster was found is inset in each panel. (H) Average motor activity overlaid onto the optic flow stimulus. (I : top) Response of probe responsive cells identified using cross correlation with stimulus regressor. (I : bottom) Response of the same cells shown on top to 1cm/s optic flow. (J) Number of cells identified as probe responsive. (K) Average response of the probe responsive cells to negative probe pulses post acclimatization and when presented at random (baseline). (L) Response amplitude of the probe responsive cells to negative probe when presented randomly compared to the amplitude post acclimatization. ns: p = 0.11 by Wilcoxon signed-rank test. Traces shown in C-I, K are mean ± SEM.

We then used correlation based hierarchical clustering to identify functional GC response types (Figure 5C-G). Clustering was performed independently for each fish using activity traces from the second trial. The number of clusters per fish was set to 5 with the help of silhouette plots. The clusters were then assigned names based on the relationship of the observed response types to the optic flow stimulus. The same response types were not present across all fish but were largely overlapping (Figure 5C-G).

A substantial proportion of cells (43%) were either inactive or sparsely active in a non-stimulus selective manner (Figure 5F). 15% of all cells were strongly tuned and an additional 31% were weakly tuned to the forward optic flow stimulus respectively (Figure 5C,E). 3% of all cells were activated non-specifically by forward optic flow and exhibited a persistent elevation in calcium activity outlasting the stimulus and/or motor activity (Figure 5D). The final cluster consisting of 8% of the sampled GC population included cells that were inhibited by forward optic flow (Figure 5G). Importantly, no response type was found tuned to the weak probe stimulus, indicating that the large amplitude responses observed downstream in the PC population were not caused by GC activity.

It is possible that the clustering method failed to identify a sparse subset of GCs tuned to the negative probe stimulus. To address this, we explicitly looked for cells showing responses to the 3 pulses of the negative probe stimuli by cross correlating a short segment of the calcium trace around the probe pulses with the stimulus regressor (Figure 5I). By setting a conservative threshold correlation of 0.5, we were able to find 3 ± 3 cells (mean ± SD) per fish that were tuned to the negative probe stimulus (Figure 5I - top, J). Since responses were not trial averaged, lower thresholds than 0.5 picked up correlated imaging noise. Cells identified using this method were not activated by the 1cm/s flow stimulus (Figure 5I - bottom). However, they had comparable response amplitude to the negative probe stimulus when presented in random order in the first trial (Figure 5K,L), suggesting that this is a sensory representation of the probe stimulus. Moreover, these results strongly suggest that the signal encoding deviation from expectation arrives at the PCs through CF inputs from the IO.

### Strong population-wide encoding of stimulus expectation in PCs correlates with lower swim latency

To test whether an expectation of the forward optic flow stimulus might influence swim responses, we looked at the latency with which optomotor responses were generated. In this experiment, fish were presented with a single trial of 100 pulses of optic flow stimuli, consisting of 90 pulses of forward optic flow at 1cm/s and 10 pulses of the negative probe stimulus (Figure S6A). The probe stimulus was pseudorandomly interspersed across the experiment. As seen earlier, fish swam consistently in response to forward optic flow and almost no swims were initiated in response to the probe stimulus or during periods of no optic flow (Figure S6B). PC calcium activity was imaged simultaneously.

We first sorted cells into forward or backward tuned based on their average calcium responses to each of the stimuli. An average response profile was constructed for each cell by stitching together the average response to forward optic flow and probe (Figure 6B). Using regression, we then classified cells as either forward tuned (Figure 6B, purple) or backward tuned (Figure 6B, black). Since there were 90 pulses of forward optic flow and 10 of probe, we obtained a good classification using a threshold correlation of 0.4 as imaging noise was averaged out effectively. Out of 69 ± 14 cells (mean ± SD) from 27 fish, 18 ± 5 were backward tuned and 45 ± 11 were forward tuned.

**Figure 6:**
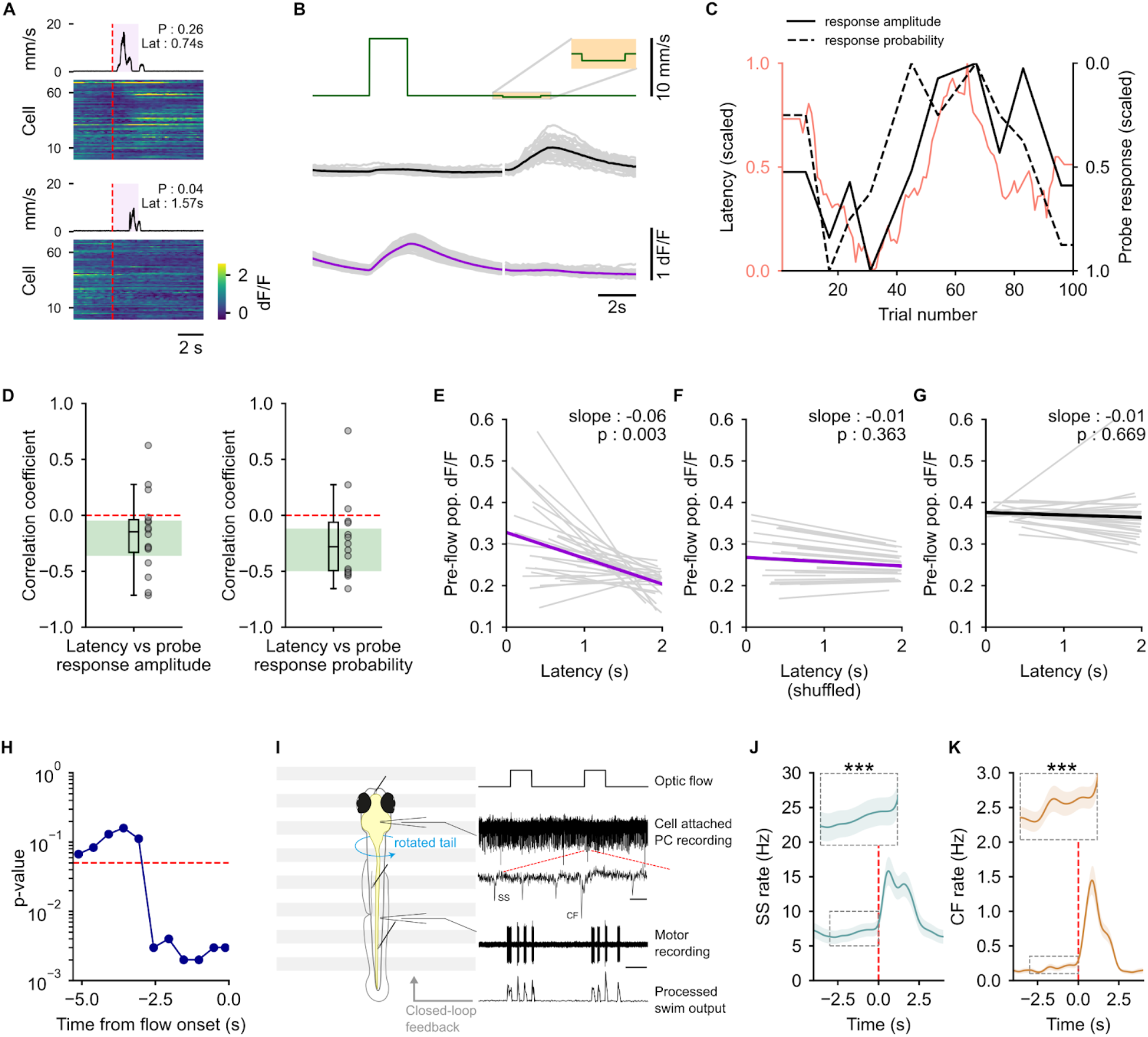
Strong population-wide representation of stimulus expectation correlates with lower swim latency. (A) Heatmap of single trial responses of all cells in a representative fish to two instances of the negative probe stimulus. The average motor activity over a window of 10 non-probe stimulus pulses around each of the probes is shown on top. The mean latency to initiate swims and the population-wide probe response probability are inset in each plot. (B) Regressor-based classification of cells into forward (magenta) and backward (black) tuned using the trial averaged responses to 1cm/s forward optic flow and the negative probe stimulus. Gray traces represent individual fish, N=27. (C) Representative plot from one fish showing latency to swim in response to forward optic flow (pink) along with probe response amplitude (black) and probability (black dashed) for backward tuned cells. (D) Both amplitude and probability are anticorrelated with latency. Green shaded area shows 95% confidence interval for the median estimated by bootstrapping. N=16 fish. (E-G) The pre-flow calcium activity is significantly anticorrelated with latency for the forward tuned cells (E). No correlation is seen when the trial order is shuffled (F) or in backward tuned cells (G). Slope and p-value by linear mixed-effects are inset in each plot. N=27 fish. (H) The pre-flow calcium activity in forward tuned cells is significantly anticorrelated with latency up to 2.5s prior to the onset of the forward optic flow stimulus. (I) Left: Schematic of the experimental preparation used for extracellular loose-patch recordings from PCs. Right: Traces showing the optic flow stimulus, PC recording with simple spikes (SS) and climbing fiber (CF) inputs, fictive motor recording and the processed motor signal used for closed-loop feedback for some fish. (J, K) Both SS (J) and CF input (K) rates show a significant, steady increase prior to the onset of optic flow. *** : p < 0.001 by linear mixed-effects. N=20 cells in (J) and N=15 cells in (K).

Next, we hypothesized that if swim responses to optic flow were influenced by an expectation based on stimulus history, a stronger expectation could lead to lower latency responses. Previous experiments have shown that an enhanced response to an unexpected probe stimulus encodes the level of deviation from expectation and can be used as a proxy for how strongly fish expect the acclimatization stimulus (Figure 4). We therefore looked at both the population averaged amplitude and the probability of occurrence of large calcium transients in response to probe stimuli in backward tuned cells - a high amplitude or probability would imply higher confidence in the expected stimulus pattern and vice versa. Response probability was estimated by counting the occurrence of a large calcium event (> 3 SDs from baseline) during probe pulses. Since probe stimuli were presented infrequently in this experiment, we linearly interpolated the amplitude and probability to obtain coarse-grained readouts of the expectation of forward optic flow across the 100 stimulus pulses (Figure 6A,C). Similarly, latency values were also linearly interpolated and smoothened with a rolling average spanning 10 stimulus pulses. Fish that paused swimming for a continuous sequence of 10 or more pulses were excluded to minimize interpolation artifacts. The 10 pulse long smoothing window matched the average distance between two probe stimuli, thereby yielding a comparably coarse-grained estimate of latency. We then proceeded to compute the Pearson’s correlation coefficient between these population-wide readouts of how strongly fish expected forward optic flow and the latency to swim. Swim latency was negatively correlated with both amplitude and response probability (Figure 6C,D). These results suggest that fish might incorporate an expectation of optic flow stimulus in planning swim responses to repetitive stimuli.

In the above experiment, if fish indeed initiated swims with lower latency because they expected the forward optic flow stimulus, there should be a neural representation of this expectation when forward optic flow stimuli were presented, whether in PCs or elsewhere. As both forward optic flow and motor activity cause strong PC activation, we decided to look at the calcium activity prior to the onset of the forward optic flow stimulus to exclude sensory and motor contributions to the dF/F signal. For this analysis, we extracted the average pre-flow dF/F for both forward and backward tuned cells and fit a linear mixed-effects model between swim latency and the pre-flow calcium activity. The pre-flow calcium activity in forward tuned cells showed a significant negative correlation with latency (Figure 6E). A significant negative correlation could be detected up to ~2.5s before the onset of the stimulus (Figure 6H). The correlation was absent when stimulus order was shuffled (Figure 6F) and was not present in cells tuned to the backward direction of optic flow (Figure 6G). This observation is consistent with the idea that the cells active during forward optic flow are likely the ones involved in the generation of behavioral responses and hence any representation of an expectation of forward optic flow that contributes to behavior is therefore also present in their pre-flow activity and not in the backward tuned cells.

We then performed loose-patch extracellular recordings from PCs in paralyzed fish presented with repetitive forward optic flow stimuli (Figure 6I). Cells that showed strong activation during forward optic flow were selected. All 20 cells that were analyzed showed a strong simple spike (SS) rate increase during forward optic flow stimuli and 15 out of the 20 showed a strong increase in climbing fiber (CF) input rate. In these cells, we saw a significant increase in both SS (Figure 6J) and CF input (Figure 6K) rate in a 3s window leading up to the onset of forward optic flow. This indicates the possible presence of a predictive component in the activity of PCs, which contributes to the initiation of swim responses to optic flow.

### Unexpected stimuli cause a cerebellum dependent transient elevation in swim latency

Next, we probed for a causal link between cerebellar activity and the expectation dependent component of the optomotor response. If expectation of a stimulus contributed to the generation of lower latency swims, we hypothesized that deviant probe stimuli would perturb behavior and lead to longer latency responses to subsequent presentations of forward optic flow. To quantify this, we looked at a window of 5 optic flow stimulus pulses before and after a probe stimulus (Figure 7A,B). The average latency in the window before the probe was considered as the baseline and a fold-change in latency from this baseline was calculated for each of the 10 stimulus pulses in the window around the probe (Figure 7B). The probe stimulus caused a significant elevation in latency in response to the next occurrence of the forward optic flow stimulus. The elevated latency returned back to baseline after a single pulse of the forward optic flow stimulus following the probe (Figure 7C). This suggests that fish indeed use expectations based on stimulus history to generate optomotor responses.

**Figure 7:**
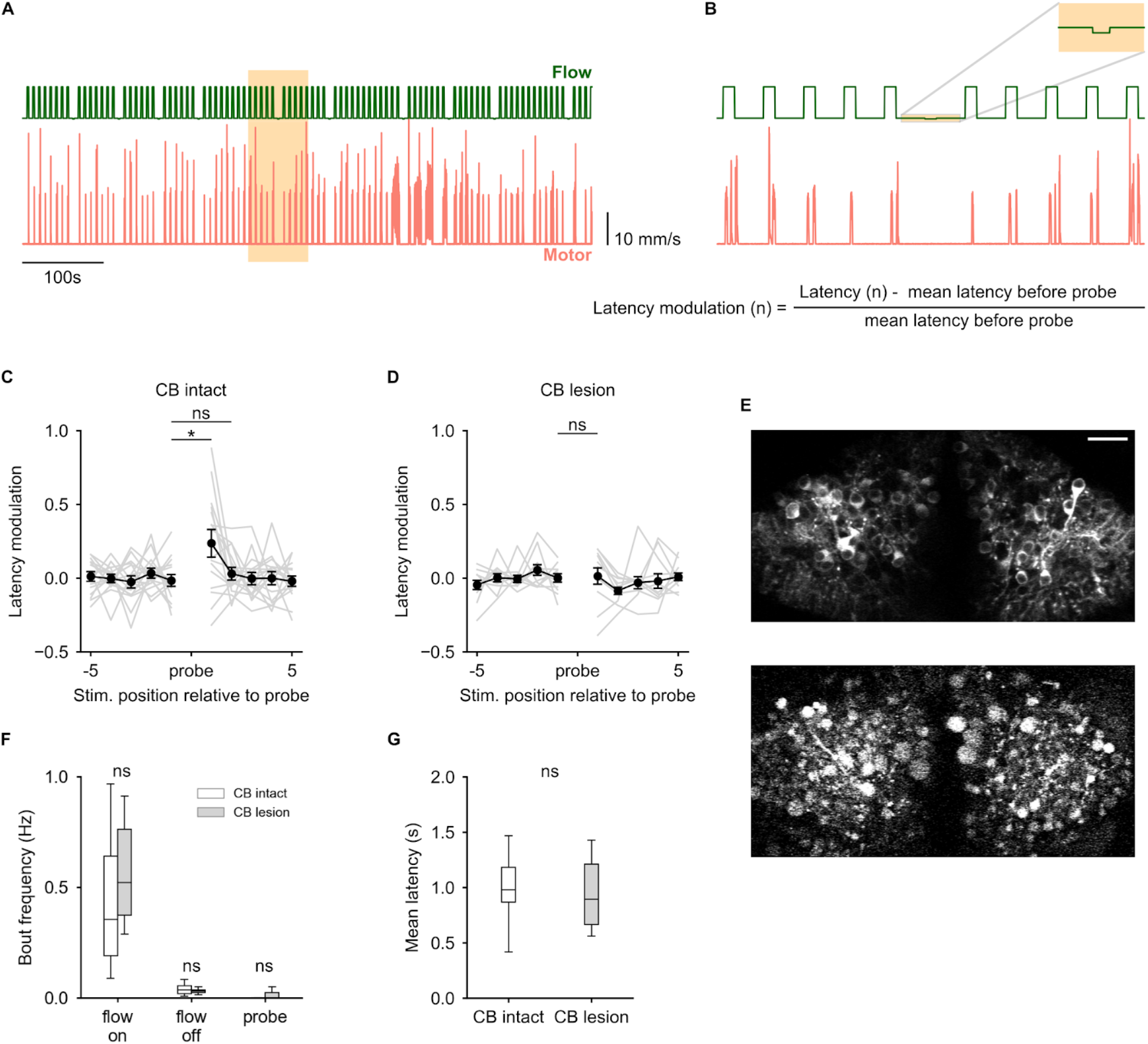
The cerebellum uses a learnt expectation of optic flow stimulus to generate swims with lower latency. (A) Optic flow stimulus protocol used (green) and corresponding motor responses of a representative fish (pink). The positions of the probe stimuli were randomized. (B) Zoomed in view of the highlighted region in (A) Top: showing a probe stimulus ± 5 forward optic flow pulses. Bottom: Calculation of latency modulation around the probe. (C) Average latency modulation for 10 forward optic flow pulses around the probe stimulus. N=17 fish, * : p = 0.03, ns : p = 0.43 by Wilcoxon signed-rank test. Unexpected probe stimuli cause a transient elevation in latency. (D) Latency modulation around the probe stimulus for cerebellum lesioned fish. N=11, ns : p = 0.79 by Wilcoxon signed-rank test. Probe stimuli did not cause an elevation in latency in lesioned fish. (E) A single two-photon optical section of the GCaMP6s expressing Purkinje cells before (top) and after (bottom) lesioning. Disintegration and blebbing of cells and neuropil can be seen after the lesioning protocol. Scale bar : 20μm. (F, G) Both lesioned and non-lesioned fish show similar levels of behavioral responsiveness (F) and respond to optic flow with similar average latency (G). ns : p > 0.05 by independent sample t-test.

In order to test a causal role for the cerebellum in this process, we lesioned the cerebellum (Figure 7E) in a different group of fish and performed the same experiment as above. Both lesioned and non-lesioned fish showed similar levels of behavioral responsiveness and the mean latency remained comparable between both groups (Figure 7F,G). However, the lesioned fish did not show an elevation in latency after the probe stimulus was presented (Figure 7D). The fact that behavior was not perturbed by an unexpected stimulus in lesioned fish indicates that the cerebellum is involved in incorporating expectations based on stimulus history in the generation of swim responses to optic flow, possibly with lower latency.

## Discussion

In this study, we investigated how the brain acquires behavioral timescale internal models of the world to respond more efficiently to predictable stimuli. Using two-photon calcium imaging in head-restrained larval zebrafish in a closed-loop optomotor environment, we identify and characterize a previously unknown expectation based component involved in the optomotor response. Our results show that with experience, fish learn to expect the direction of optic flow stimulus. PCs in the cerebellum are strongly activated by optic flow in a strictly direction selective manner (Figure 1). In addition to encoding sensory input, the PC response to optic flow included a component which encoded the degree of unexpectedness of the stimulus, while maintaining direction tuning (Figures 2, 4). As seen in other systems ^4,16,33^, we propose that this non-sensory component constitutes an error signal which is critical for maintaining an updated belief of what to expect from the sensory world. Activity in the GC population revealed a variety of sensory tuned responses but not the error signal, indicating that CF inputs from the IO convey error information to PCs (Figure 5). A subset of GCs exhibited a persistent stimulus tuned response, which possibly encodes stimulus expectation (Figure 5D,G). An error driven learning rule could enable PCs to incorporate appropriate predictive signals from GCs to modulate behavior. The PC population-wide encoding of error, which is a proxy for how strongly fish expect a particular stimulus, correlated negatively with swim latency, suggesting that stimulus expectation might indeed contribute to lowering the latency of swim responses (Figure 6A-C). Subsequently, through lesion experiments, we identify a causal role for the cerebellum in latency modulation when unexpected stimuli are encountered (Figure 7). In sum, these results point to a central role for the cerebellum in signaling deviations from expectation and in sculpting motor responses for expected and unexpected stimuli.

### Role of the cerebellum in zebrafish optomotor behavior

Previous studies have identified the larval zebrafish pretectum as the primary optic flow processing region involved in the optomotor response ^25,34^. The cerebellum receives inputs from direction tuned neurons in the pretectum ^34^ but a direct role for the cerebellum in sensorimotor transformations involved in the optomotor response has not been identified ^25,34^. Here we show that lesioning the cerebellum does not affect the response latency or swim probability (Figure 7F,G). However, unlike control fish, the lesioned fish were not perturbed by an unexpected change in optic flow direction (Figure 7C,D), suggesting an inability to incorporate an expectation of optic flow direction into the optomotor response. This shows that the cerebellum does not play a direct role in the generation of the response but is rather involved in biasing behavior based on past experience.

### Possible circuit mechanism

The findings presented here give us important insights into the mechanisms employed by the cerebellar circuitry to incorporate an internal model of the sensory world into behavior generation. These are discussed below.

### Dense and persistent sensory representations in GCs

The 7dpf larval zebrafish cerebellum consists of around 6000 GCs which converge onto 300 PCs. Our observation of dense activation of the GC population in response to optic flow stimuli (Figure 5C-G) is consistent with recent findings from larval zebrafish ^35,36^ and other species ^3,37^. We also found response profiles in the GC population exhibiting stimulus tuned but persistent elevation in calcium activity (Figure 5D,G). One of these response profiles, present in 6 out of 8 fish (Figure 5G), was inhibited strongly by forward optic flow and showed long lasting elevation during periods of no optic flow or probe. It is possible that sensory expectations could be encoded in persistent activity of such a subpopulation. A recent study of GC responses to luminance changes in larval zebrafish also found temporally diverse and persistent activity lasting tens of seconds from stimulus onset ^36^. Depression of GC inputs coincident with error signals from the IO would allow PCs to select response profiles that contribute to error free behavior ^6,38^.

### Error signals are encoded in CF inputs from the IO

No error representation was present in the GC calcium responses, even though a substantial proportion of them were sampled (Figure 5). This strongly suggests that the error signal we see in the PC population, representing the deviation from expected direction of optic flow, is encoded in CF inputs from the IO. Furthermore, it has been observed that large calcium signals recorded from PCs are more likely to be caused by CF driven bursts or calcium spikes than changes in simple spike rates caused by GC activity. It is therefore possible that PC direction tuning (Figure 1E) determined by calcium imaging reveals how error signals are tuned for each cell rather than their sensory tuning. Future experiments to determine how error maps onto GC input will provide important insights into cerebellar computation.

How the brain generates error signals has been a long standing question in neuroscience. In order to generate error, there must already be a prediction or an expectation which is compared with ground truth sensory input to determine whether there are mismatches. This error can then be used to update the expectation such that future errors are minimized. In the findings presented here, the error signal is most likely computed by or relayed to IO neurons, which then map onto PCs in the cerebellum. This would indicate that neurons encoding the expected direction of optic flow, either as persistent activity or distributed across the network, are present upstream of the cerebellum.

### Integration of sensory expectations and error by PCs and their influence on behavior

Individual PCs receive convergent input from a large number of GCs, possibly composed of a variety of different response types. We hypothesize that PCs implement an error driven learning rule that adaptively scales the contribution of GC response types to behavior. Specifically, the inputs to PCs from GCs encoding stimulus expectation that are active during probe get progressively suppressed, thereby contributing less to generating the behavioral response. Multiple observations presented here are consistent with this model. (i) The expectation is updated only when stimuli that are either consistent (repeated forward flow) or inconsistent (deviant probe flow) with the past are experienced (Figure 4). Extended periods of no optic flow do not lead to a change in the amplitude of the error response (Figures 4G, S3), nor does constant optic flow (Figure S4). These results point towards an experience gated updating process. (ii) The latency to swim in response to forward optic flow is negatively correlated with the prevalence of error encoding in the PC population (Figure 6A-C). Furthermore, the response latency is longer immediately following a probe pulse compared to the pre-probe baseline (Figure 7C) but no elevation is seen in cerebellum lesioned fish (Figure 7D). This suggests that stimulus expectation might indeed contribute to a lower latency optomotor response.

While the model proposed here provides an algorithmic description of the neural mechanism, it opens up several questions for further investigation. As discussed above, for an error signal to be computed, a comparison between an expectation and ground truth is necessary. It is unclear how the brain acquires this expectation and how it is encoded in neural activity. Examining the activity of IO neurons and the inputs they receive will shed light on this process. Additionally, the role of the cerebellum may not be limited to tuning the influence of expectation on behavior generation. Cerebellar feedback to upstream brain regions could play an important role in acquiring these expectations. Lastly, the output of the cerebellum does not directly cause movements. It remains to be seen how cerebellar output integrates with sensorimotor computations involved in generating the optomotor response.

### Ethological relevance of predicting optic flow

Shallow water streams inhabited by larval zebrafish often show repetitive and predictable changes in flow due to water waves or vortex streets shed by objects intercepting laminar flow. Fast and precise reactions to these perturbations caused by flowing water are likely to prevent fish from being washed away from a favorable environmental niche. In our experiments, fish learnt to expect optic flow direction with minimal prior exposure and continuously adapted their internal model with experience. Such an adaptable predictive mechanism functioning on timescales of immediate behavioral requirements is highly relevant in the natural setting. In addition to this homeostatic function, it is possible that fish incorporate predicted environmental perturbations to fine tune other behavioral responses such as escape swims or prey capture maneuvers, such that the motor command generated in these situations compensates for displacement due to expected flow patterns. This study sets up an ideal model to investigate how multiple parallel predictive mechanisms involved in expecting flow induced perturbations and those involved in behaviors such as escape or prey capture, work in concert to ultimately guide appropriate behavioral choices.

### Conclusion

Using the accessibility of the larval zebrafish model, we showed that Purkinje cells in the cerebellum can encode expectations of sensory stimuli and report error when the expectation is not met. Such cerebellar computations are incorporated into motor planning so as to generate swims with quick reaction times. These results point to important and conserved functions of the cerebellum in acquiring and updating internal models of the world.

## Methods

### Fish

We used 6-8dpf zebrafish (*Danio rerio*) larvae for all experiments. The larvae were raised in E3 medium (composition in mM: 5 NaCl, 0.17 KCl, 0.33 CaCl_2_, 0.33 MgSO_4_, 0.5 methylene blue) or fish facility water with 0.5mM methylene blue, at 28.5°C in accordance with standard protocols (https://zebrafish.org/home/guide.php). Tg(aldoca:GCaMP6s);nacre^-/-^ larvae (from Prof. Masahiko Hibi, Nagoya University, Japan) and Tg(neurod:GCaMP6F);nacre^-/-^ larvae (from Prof. Claire Wyart, I.C.M., Paris) ^39^, outcrossed into the Indian wildtype (from local aquarium suppliers) background were used for experiments involving two-photon imaging. Indian wildtype larvae were used for electrophysiological recordings. Each experiment was spread out over at least 3 different batches of larvae. All experimental procedures were approved by the Institutional Animal Ethics Committee of the National Centre for Biological Sciences.

### Experimental preparation

For experiments involving two-photon imaging, 7-8dpf zebrafish larvae were anesthetized by briefly immersing them in 0.017% MS222 (Sigma-Aldrich) and embedded dorsal side up in a drop of 2% low gelling temperature agarose (Sigma-Aldrich) placed in the center of a 60mm petri dish lid. The dish was filled with E3 medium and the tail below the level of the swim bladder was freed from agarose. The embedded fish were allowed to recover for 4-8 hours before the experiment.

### Real-time motor output detection for closed-loop optomotor feedback

Custom Bonsai workflows ^40^ were used for real-time motion detection. The algorithm used by our software for online motion detection closely matched previously published protocols ^28,41^. Video based motion detection was done at 200fps using a Flea3-USB3.0 camera (FLIR). The tail of the fish was illuminated from one side by a collimated 780nm LED (M780D2, Thorlabs) and imaged onto the camera sensor below (Figure 1Aa), using a reversed Nikon 50mm f/1.8d lens with a 24mm extension tube for increased magnification. Both the illumination and imaging light paths were filtered through 780nm bandpass filters (FBH-780-10, Thorlabs). The imaging light path was filtered to exclude light from the visual stimulus and two-photon laser and the illumination light path was filtered to block low intensity visible light emission from the 780nm LED (Figure 1A). Real-time motion detection was performed by frame subtraction. The fraction of motion pixels in each frame, which scales with the amount of tail movement, was lowpass filtered with a 10ms time constant to obtain the instantaneous swim strength (ISS). This lowpass filter simulates the inertial effects experienced by larval fish in water. The ISS was streamed in real-time using Open Sound Control (OSC) protocol to custom software written in Processing 2.2.1 (https://processing.org/). This software generated the visual stimulus, executed the experimental protocol, and generated TTL signals to synchronize two-photon imaging or electrophysiology data acquisition with visual stimulus presentation.

ISS for the electrophysiology preparation was estimated from fictive swims recorded unilaterally from the ventral root (Figure 6I). Analog output from the recording amplifier was digitized at 12KHz in 10ms chunks (PCIe-6361, National Instruments) and acquired using a Bonsai workflow. Each 10ms chunk was bandpass filtered between 0.2-5kHz. The standard deviation of each chunk was calculated and the offset induced by recording noise was subtracted. This signal was lowpass filtered at 10Hz and thresholded manually to obtain an estimate of the ISS, which was streamed to the experiment control software.

The ISS was scaled individually for each fish to obtain a realistic estimate of swim velocity. 1-3 trials of optic flow at 10mm/s lasting 10-15s were presented before the start of each experiment in the open-loop configuration. A scaling factor, *s*, was calculated from these trials such that the ISS when scaled resulted in an average swim bout velocity of 12-15mm/s. This process is described by the following equation.

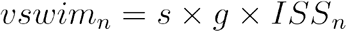

Where *n* is the frame count. *vswim* is the instantaneous velocity of the fish in the closed-loop environment. *s* is the scaling factor set individually for each fish. *g* is the gain of the closed-loop feedback, set at 1.0 for all experiments. *ISS* is the instantaneous swim strength.

### Visual stimulus

Custom software written in Processing 2.2.1 was used to control the experimental protocol as well as to generate and update the visual stimulus at 120Hz. The visual stimulus consisted of square wave grating with a spatial period of 10mm presented 5mm beneath the fish. The grating velocity was a combination of both the stimulus velocity and the scaled *ISS*, as per the following equation.

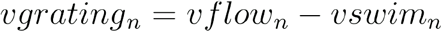

Where *vgrating_n_, vflow_n_* and *vswim_n_* are the instantaneous velocities at frame *n* of the grating stimulus, externally imposed optic flow and the fish respectively.

In the calcium imaging setup, visual stimulus was projected on a 60mm diameter diffusive paper screen placed 5mm below the fish. The projector was custom built around a 5-inch LCD panel (Adafruit) removed from its housing. The LCD was illuminated from behind with a collimated 617nm LED (Luxeonstar) and was imaged onto the projection screen. A custom made driver was used to power the LED antiphase to the linescans of the fast x-galvo during two-photon imaging. This eliminated the possibility of bleedthrough of light from the visual stimulus into the calcium imaging channel. In addition to this, a longpass filter (575lp, Chroma) was placed in the projection light path to prevent harmful levels of light from reaching the PMT. The final pixel size on the projection screen was 0.32mm. A 5X5mm window was cut out in the projection screen to facilitate video recording of tail movements from below (Figure 1A). Visual stimulus for the electrophysiology experiments was presented directly on an LCD screen positioned beneath the fish.

### Experimental protocols

#### Optic flow stimuli

Optic flow stimuli were presented in 2s long pulses, separated by an interval of 5s with stationary gratings. Each trial consisted of multiple pulses of optic flow stimuli and trials began with 5s of stationary gratings followed by a 2s pulse. During this 2s pulse, the gratings moved continuously, in each frame, for the 1cm/s stimuli. In case of the probe stimuli, the gratings moved in 1-pixel steps a total of 4 times (Figures 1–4) or 1 to 3 times (Figures 5, 6) during the 2s period. These stimuli are represented in the figures by their average velocity. In experiments involving more than one trial, an inter-trial interval of 30s was introduced between trials when stationary gratings were presented. Imaging began at the onset of every trial and was stopped 6s after the trial ended, resulting in a 24s gap in two-photon acquisition between trials. Behavior data was acquired continuously throughout the whole protocol.

#### Two-photon calcium imaging

Imaging was performed on a custom two-photon microscope controlled using ScanImage 3.8 ^42^. A Ti-Sapphire laser (Chameleon Ultra II, Coherent) tuned to 920nm was used to excite GCaMP6s. The laser was focussed at the sample using a 20X 1.0NA objective lens (XLUMPLFLN20XW, Olympus), downstream of 50mm and 300mm focal length scan and tube lenses (AC-300-050B and AC-508-300-B, Thorlabs). Scanning was achieved using 3mm aperture scan mirrors (6215H, Cambridge Technology). Fluorescence emission from GCaMP was bandpass filtered (FF01-520/70, Semrock) and detected with a GaAsP PMT (H7422PA-40, Hamamatsu). Output from the PMT was amplified using a preamplifier (SR570, Stanford Research Systems) and acquired using the same data acquisition card used to drive the scan mirrors (PCI-6110, National Instruments). Data acquisition from the microscope was triggered by a TTL pulse generated at the beginning of each trial by an arduino connected to a separate computer running the behavior protocol. A continuous timeseries of two-photon images was acquired for every trial of the experimental protocol. Multi-trial protocols had an inter-trial interval of 30s. Imaging was discontinued for 24s out of the 30s interval.

#### Purkinje cell calcium imaging

Head-restrained Tg(aldoca:GCaMP6s);nacre^-/-^ larvae were used to image PC calcium activity. A single 512X128 pixel field of view with 0.53μm pixel size, centered around the Purkinje cell layer was scanned using a pixel dwell time of either 1.6μs (Figure 6) or 3.2μs (all other experiments), resulting in frame rates of either 7.8Hz or 3.9Hz respectively. Laser power measured at the sample plane was 10-15mW for 3.2μs pixel dwell time and 20-30mW when the pixel time was set at 1.6μs.

#### Granule cell calcium imaging

Head-restrained Tg(neurod:GCaMP6f);nacre^-/-^ larvae were used to image GC calcium activity. 6 planes separated by 10μm were imaged in every fish, starting from the dorsal side of the GC layer. The two-trial behavior protocol was carried out serially for each imaging plane. Since galvo scanning was used, we scanned a single hemisphere of the GC layer to achieve sufficient frame rate for GCaMP6f, while still densely sampling the cell population. A 256X128 pixel field of view with 0.53μm pixel size was scanned with a pixel time of 3.2μs, resulting in a frame rate of 7.8Hz. Laser power used for GC imaging was in the range of 15-20mW at the sample plane. Post acquisition, a 2-frame temporal averaging was used to improve signal to noise ratio to aid subsequent analysis.

#### Two-photon laser ablations

Tg(aldoca:GCaMP6s);nacre^-/-^ larvae were prepared for cerebellum ablations using the same protocol as used for the calcium imaging experiments. 10 planes within the Purkinje cell layer, separated by 4μm, were serially scanned with a pixel dwell time of 1.6μs. Ablations were confined to the Purkinje cell layer using visual aid from GCaMP fluorescence. The laser was tuned to 920nm with a power of ~230mW at the sample plane. Each plane was scanned 10 times before moving to the next. This led to effective plasma mediated lesioning of the whole cerebellar region being scanned. Lesions were confirmed visually by an immediate increase in fluorescence intensity due to uncontrolled influx of calcium into the cytosol, followed by blebbing and disintegration of cells. Fish were allowed to recover for 1 hour post lesion before initiating the behavior protocol. The cerebellum was scanned while the behavior protocol was carried out, with laser power and scan settings identical to those used for calcium imaging in non-lesioned fish to maintain experimental conditions.

#### Electrophysiology

6-7dpf zebrafish larvae were first anesthetized in 0.01% MS-222 and transferred to a recording chamber. Two pieces of fine tungsten wire (California Fine Wire) were used to pin the larvae through the notochord onto a Sylgard (Dow Corning) slab in the recording chamber. A third pin was used to orient the head dorsal side up (Figure 6A). The MS-222 was then replaced by external solution (composition in mM: 134 NaCl, 2.9 KCl, 1.2 MgCl_2_, 10 HEPES, 10 glucose, 2.1 CaCl_2_, 0.01 d-tubocurarine; pH 7.8; 290 mOsm) and the skin covering the brain and along the tail was carefully removed using forceps (Fine Science Tools). The preparation was then taken to a rig apparatus.

Patch pipettes for loose patch recordings were made using thick-walled borosilicate capillaries (1.5 mm OD; 0.86 mm ID, Warner Instruments) and pulled to 1-1.5 μm tip diameter using a Flaming Brown P-97 pipette puller (Sutter Instruments), such that their resistance when backfilled with external solution was approximately 7-10 MΩ. Similarly, suction recording pipettes for fictive motor recordings were made using thin-walled borosilicate capillaries (1.5mm OD; 1.1mm ID, Sutter Instruments), pulled to ~5 μm tip diameter, such that their resistance when backfilled with external solution was ~0.7-1.5 MΩ.

First, ventral root recordings to monitor fictive swimming were obtained by guiding the pipette to a myotomal boundary (~15-20th somite) and applying mild suction. Once a successful recording was obtained, the focus was shifted to the cerebellum and loose patch recordings from Purkinje neurons were obtained. The pipettes were guided visually using a 60x/1.0NA water immersion objective of a compound microscope (Ni-E, Nikon) and a motorized micromanipulator (PatchStar, Scientifica). Once both recordings were obtained, the condenser of the microscope was gently moved to allow the introduction of a 5-inch LCD screen (Adafruit) to present visual stimuli.

Visual stimulus presentation and electrophysiological recordings were triggered simultaneously and acquired using a Multiclamp 700B amplifier, Digidata 1440A digitizer and pCLAMP software (Molecular Devices). The data were low-pass filtered at 2kHz using a Bessel filter and sampled at 50kHz at a gain between 10-2000, adjusted depending on the quality of each recording. The ventral root recordings were parallely acquired and real-time analysis was performed to provide closed-loop optomotor feedback as described above.

#### Calcium trace extraction

Calcium traces were extracted from individual Purkinje cell somata using a semi-automated processing pipeline written in Python. The images were corrected for translation along the X-Y plane using phase correlation with the registration module in scikit-image ^43^. For multi-trial experiments, all frames were aligned to the first frame of the first trial and the accuracy of the registration was verified manually for every trial.

ROIs marking individual cells were drawn using a semi-automated process. For Purkinje cells, the location of individual cells was identified manually using a GUI. A three layer neural network trained using a subset of data from these experiments was used to mark cell boundaries, guided by the manually identified coordinates. For granule cells, coordinates were identified using a 2D local minimum intensity function, followed by manual pruning. Cell boundaries were then drawn by detecting bright cytoplasmic GCaMP fluorescence using a threshold function applied radially around the coordinates identified. Overlapping pixels assigned to more than one ROI were excluded from any of the assigned ROIs to avoid signal contamination from neighboring cells.

Pixels within the ROI were averaged to obtain fluorescence intensity of each cell for a given frame. The fold-change in fluorescence intensity over baseline (dF/F) was calculated with the baseline estimated as the tenth percentile of the fluorescence intensity for each cell over time. In case of multi-trial experiments, the baseline was updated once for each trial. For the continuous 700s long recordings shown in Figure 6, baseline was calculated for each frame as the tenth percentile of a one minute window around that frame. 2-frame temporal averaging was used for all analyses in case of granule cells to improve signal to noise ratio. A 2-frame temporal averaging was performed for the regression analysis for Purkinje cells.

#### Data processing and analysis

Data processing and analysis was done primarily using Python (python.org). Event detection for the electrophysiology recordings was done using MATLAB (Mathworks).

#### Behavior data processing

Swim velocity data and timestamps saved from the experiment control software at ~120Hz were used for estimating latency. A simple threshold was sufficient to detect the beginning and end of swim bouts as the signal to noise ratios were sufficiently high.

#### Estimating latency modulation

Latency modulation around the probe stimulus was estimated by determining the fold change in response latency relative to baseline latency (Figure 7A,B). The baseline was determined as the mean latency observed across 5 pulses of 1cm/s forward optic flow before the probe pulse. This normalization was necessary as slow drifts in latency developed over the duration of the experiment. Traces were selected if there were at least 3 swim bouts on either side of the probe stimulus within a window of 5 forward optic flow stimuli. A minimum of 3 such traces were required per fish to obtain an average latency modulation trace for that fish. These criteria were pre-selected before the analysis was performed and the statistical inference did not change when we later varied the selection criteria to include probe stimuli with at least 1 swim bout on either side of the probe. 17 out of 27 non-lesioned fish (Figure 7C) and 11 out of 11 cerebellum lesioned fish (Figure 7D) passed these exclusion criteria.

#### Spike detection and analysis

Spikes were identified as events that showed large fluctuations relative to a 25ms rolling window baseline and the peak times and amplitudes were extracted. Small amplitude events were classified as simple spikes (SSs) and large amplitude events as climbing fiber (CF) inputs ^21^. Once identified and sorted, recordings were triggered to the start of optic flow trials or probe trials. Mean firing rates for optic flow and probe trials were estimated by convolving peri-event time histograms of 20ms binwidth with a 200ms gaussian kernel. Analysis was done using custom scripts written in Python. CF/SS event detection was done in MATLAB (https://github.com/wagenadl/mbl-nsb-toolbox).

#### Statistics

All statistical tests were performed in Python using SciPy ^44^, statsmodels ^45^ and scikit-posthocs ^46^. The statistical tests used and their results are mentioned in the figure legends. In cases where linear mixed-effects is used, both random intercept and random slopes were included to test whether the population showed a non-zero slope. Fitting was done using the formula interface in the statsmodels package.

## Acknowledgements

The authors would like to thank the following sources of funding support: Wellcome Trust-DBT India Alliance Intermediate and Senior fellowships (VT), Department of Biotechnology (VT), Science and Engineering Research Board, Department of Science and Technology (VT), Department of Atomic Energy (VT), NCBS-TIFR graduate student fellowship (SN and AV). The authors would also like to thank Prof. Masahiko Hibi for providing the Tg(aldoca:GCaMP6s);nacre-/- fish line and Dr. Claire Wyart and Dr. Andrew Prendergast for the Tg(neurod:GCaMP6f);nacre-/- fish line. Further thanks are also due to Mr. T.P. Jagadeesh for the maintenance of our fish lines and to Drs. Lena Robra, Mehrab Modi and Ebi George for providing advice and assistance. In addition, we would like to thank the Central Imaging and Flow Facility at NCBS for support.

---

**Figure S1:**
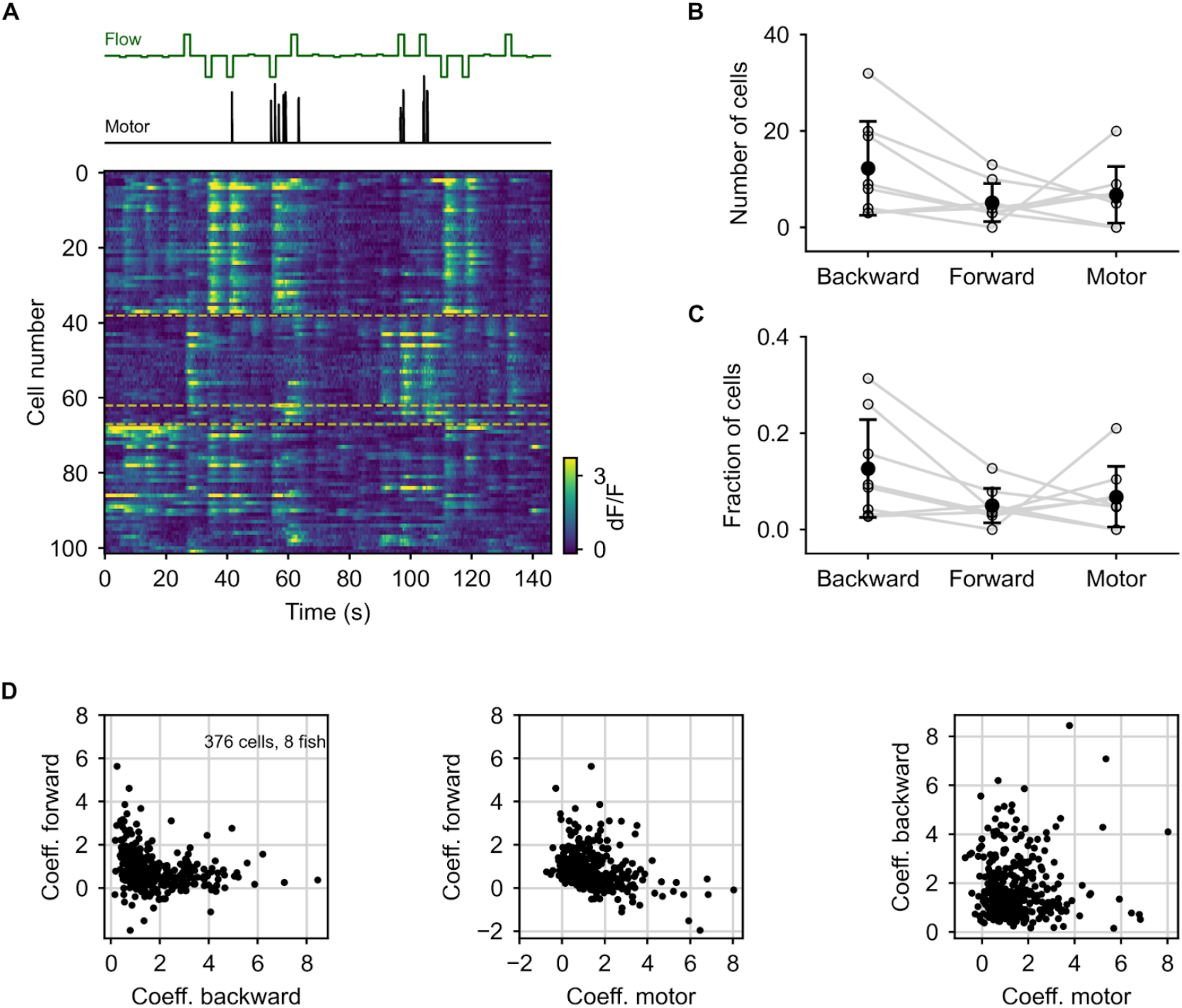
Regression-based categorization of PC response tuning. (A) Heat map of calcium activity of all cells imaged in a representative fish in response to a randomized optic flow stimulus sequence. The optic flow stimulus and motor activity of the fish are shown above. Cells are sorted based on the highest regression coefficient as backward tuned (cells 0-37), forward tuned (cells 38-62) or motor tuned (cells 63-68). None of the cells showed strong responses to both forward and backward optic flow. Cells within each group that had a strong individual correlation (Pearson’s r ≥ 0.6) with the respective regressor were then picked. (B, C) The total number of cells in each group (B) and the fraction of all cells that could be classified (C). Gray lines represent individual fish, error bars show SD. (D) Regression coefficients for all cells that could be classified. Cells responding strongly to forward optic flow did not show strong responses to backward optic flow and vice versa (A, D-left panel).

**Figure S2:**
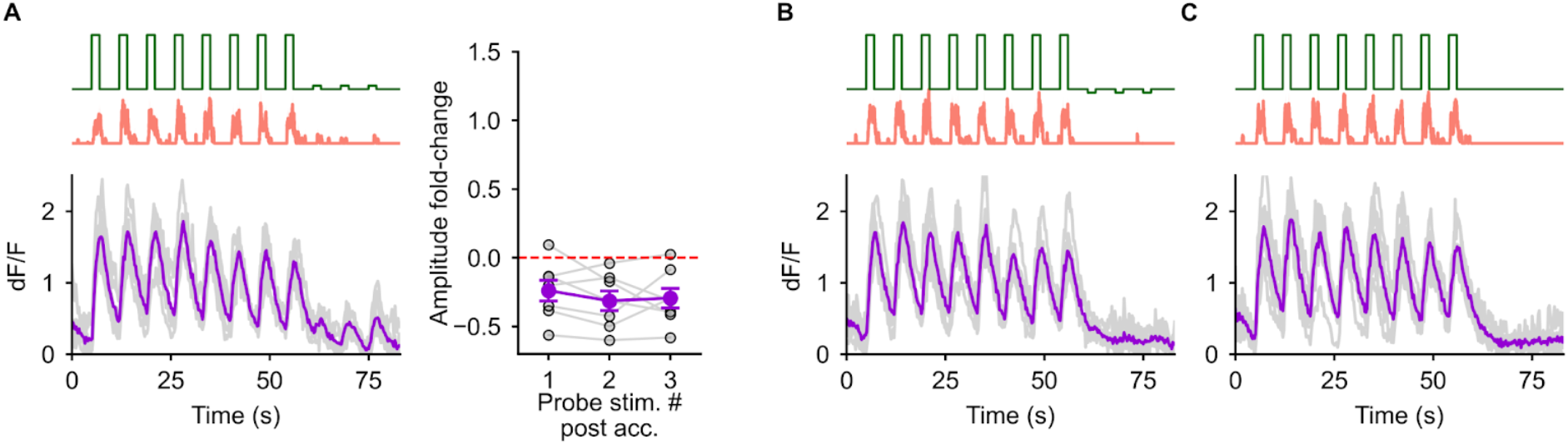
Forward tuned cells do not show an enhanced response to positive probe. (A) Left: Calcium response of the forward tuned cells to the positive probe trials (N=7 fish). Right: fold-change in amplitude of the response to positive probe stimuli post acclimatization w.r.t baseline. No increase in probe response amplitude can be seen post acclimatization. (B, C) Calcium response of the forward tuned cells in the negative probe trials (B) and in trials with no probe stimulus after acclimatization (C). No response is seen in forward tuned cells in either of these trials during the probe phase.

**Figure S3:**
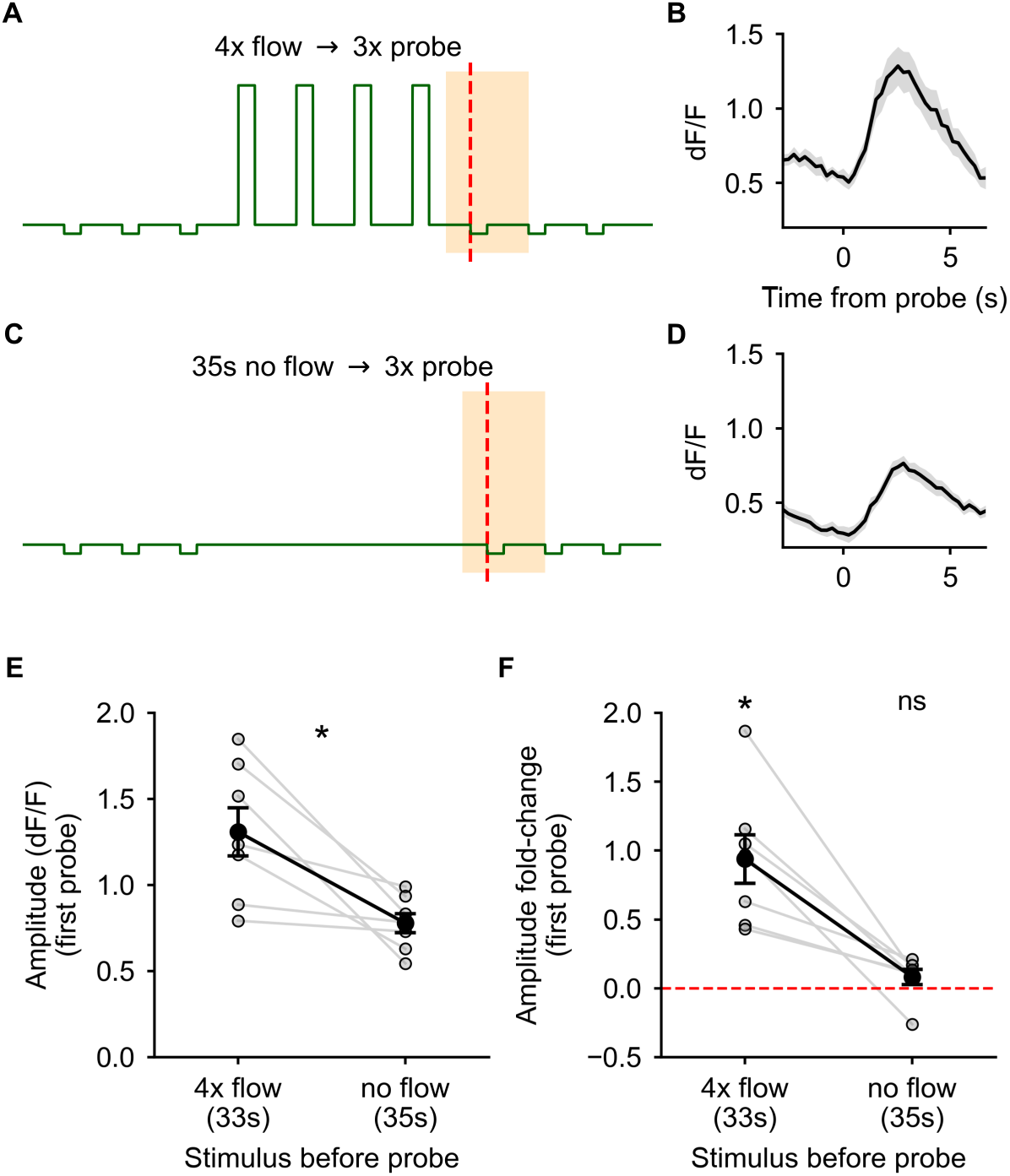
No enhancement of the response to probe stimulus is seen after extended periods without optic flow. (A, B) The optic flow stimulus in trials with 4 acclimatization pulses (A) and the average response of the backward tuned cells to the first post-probe, aligned to the onset of the probe stimulus (highlighted region in A). Shaded region is SEM across N=8 fish. (C) The optic flow stimulus between two trials showing the 35s interval with no optic flow between the end of the last probe pulse in the previous trial and the first probe pulse in the next trial. (D) The average calcium response of the same backward tuned cells to the first probe stimulus (highlighted region) after the 35s interval with no optic flow. (E) The peak amplitude of the response to probe after 4 acclimatization pulses is significantly greater than the peak amplitude after no optic flow for an equivalent time duration. * : p = 0.016 by Wilcoxon signed-rank test. (F) The amplitude post acclimatization is significantly greater than the baseline (zero) whereas the amplitude after the interval with no optic flow is comparable to the baseline amplitude. * : p = 0.016, ns : p = 0.3 by Wilcoxon signed-rank test.

**Figure S4:**
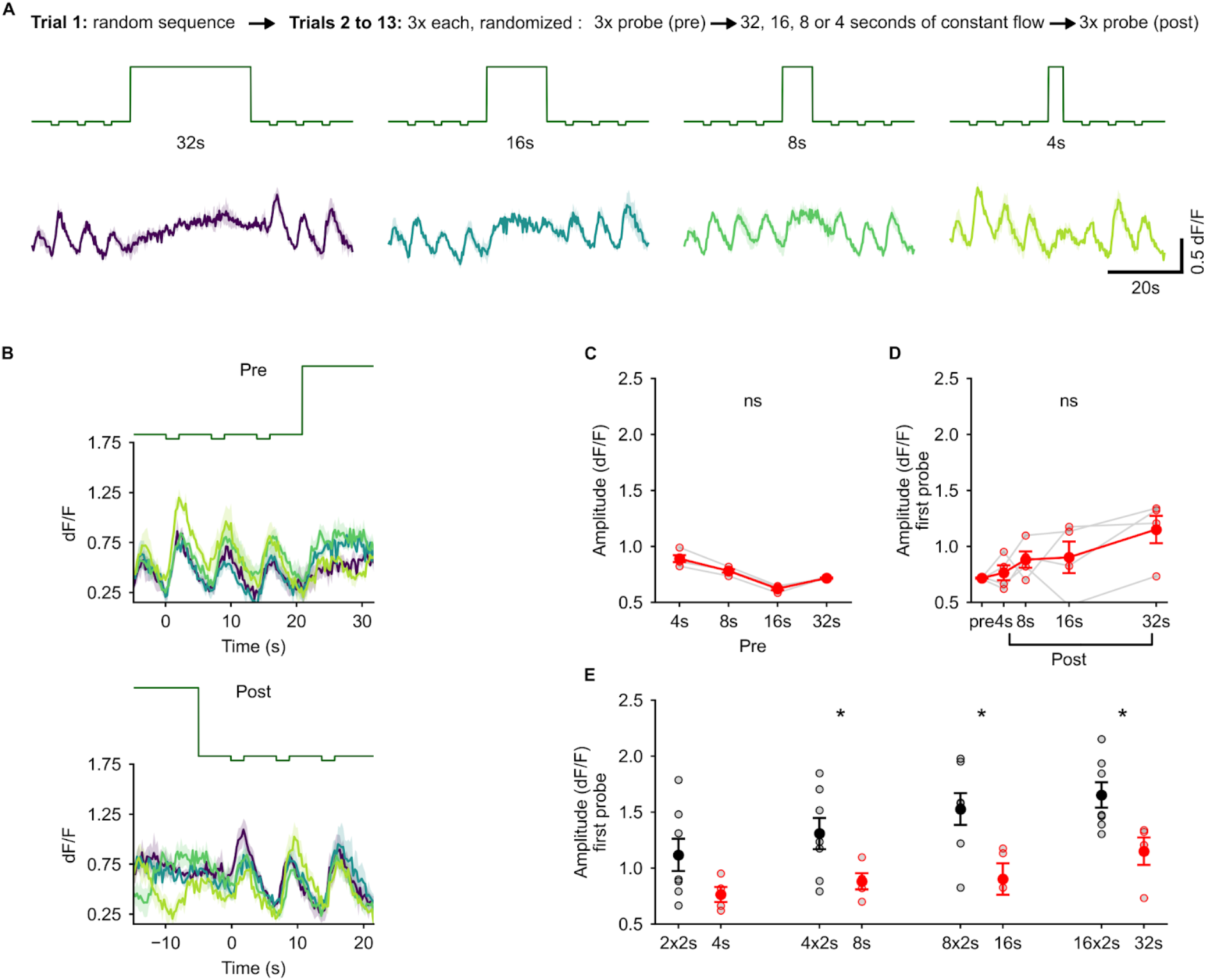
Acclimatization to constant optic flow does not lead to enhanced responses to probe. (A) Top: description of the experimental protocol. Middle (green): Stimuli presented in this experiment. A constant 1cm/s forward optic flow stimulus is presented in the acclimatization phase for 32s, 16s, 8s or 4s, corresponding to an equivalent flow-on time duration as the pulsed acclimatization stimuli in Figure 4A. Bottom: Average responses of backward tuned cells to the stimulus shown above. Shaded regions represent SEM across N=4 fish. Colors indicate the duration of the acclimatization stimulus. (B) Zoomed in view of the calcium traces shown in (A), aligned to the onset of the first pre-probe stimulus (top) and to the onset of the first post-probe stimulus (bottom). (C, D) No significant scaling of the amplitude of the pre- or post-probe stimuli could be seen. ns : p = 0.16 (C) and p = 0.08 (D) by Friedman test. (E) Response amplitudes for the first probe stimulus post acclimatization with the pulsed (black, N=7 fish) vs constant optic flow (red, N=4 fish) of equivalent durations. The probe response amplitudes after pulsed acclimatization are significantly greater than after acclimatization to constant optic flow. * : p < 0.05 by Wilcoxon rank-sum test.

**Figure S5:**
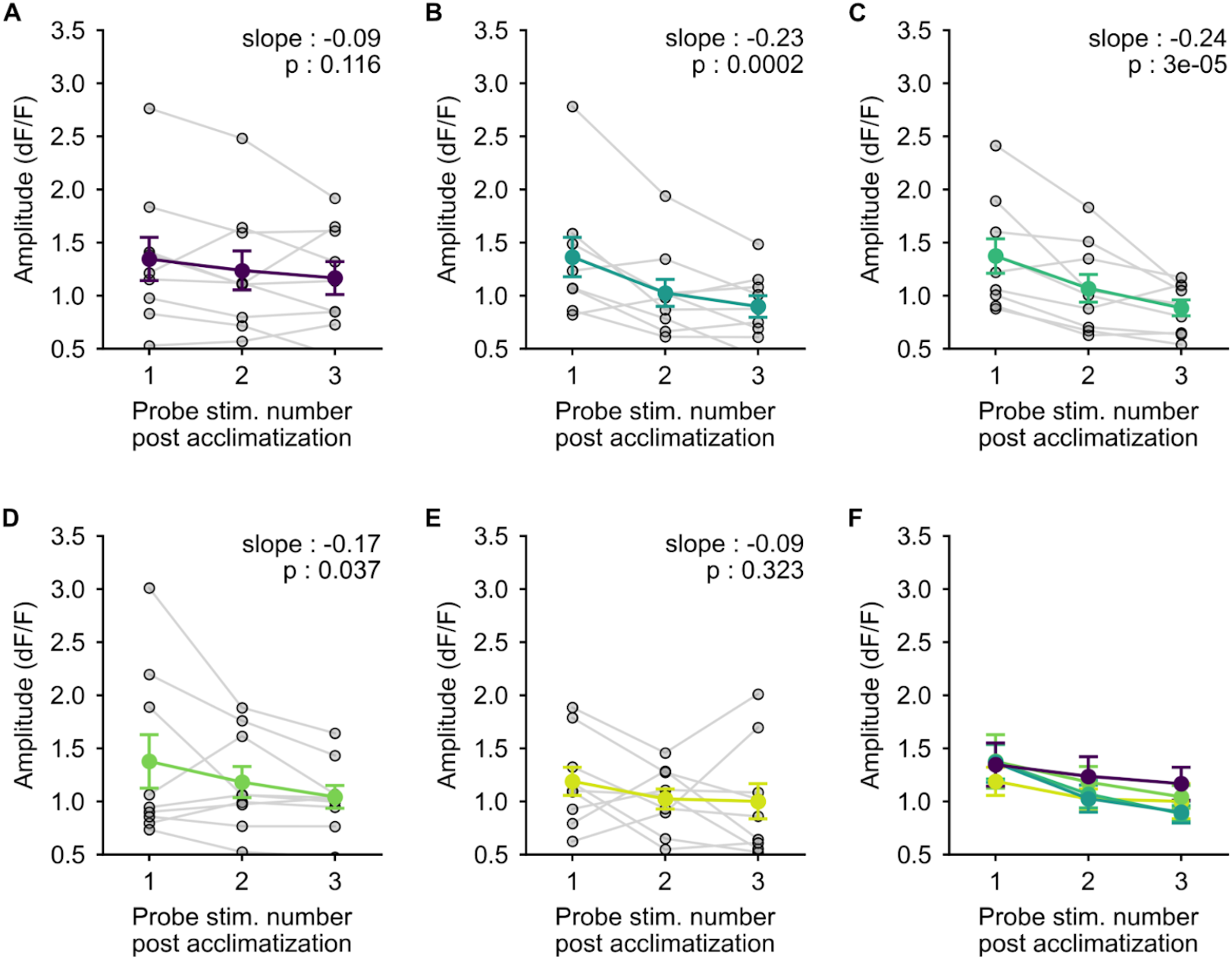
Repeated presentations of the post-probe stimulus causes a decay of acclimatization induced enhancement of the probe response. (A-E) Decay of the amplitude of the enhanced response to the post-probe stimulus over the three successive post-probe pulses for each of the delays starting from 15s (A) to 1s (E). Slope and p-value by linear mixed-effects are inset in each plot. (F) Average amplitude values from (A-E) overlaid.

**Figure S6:**
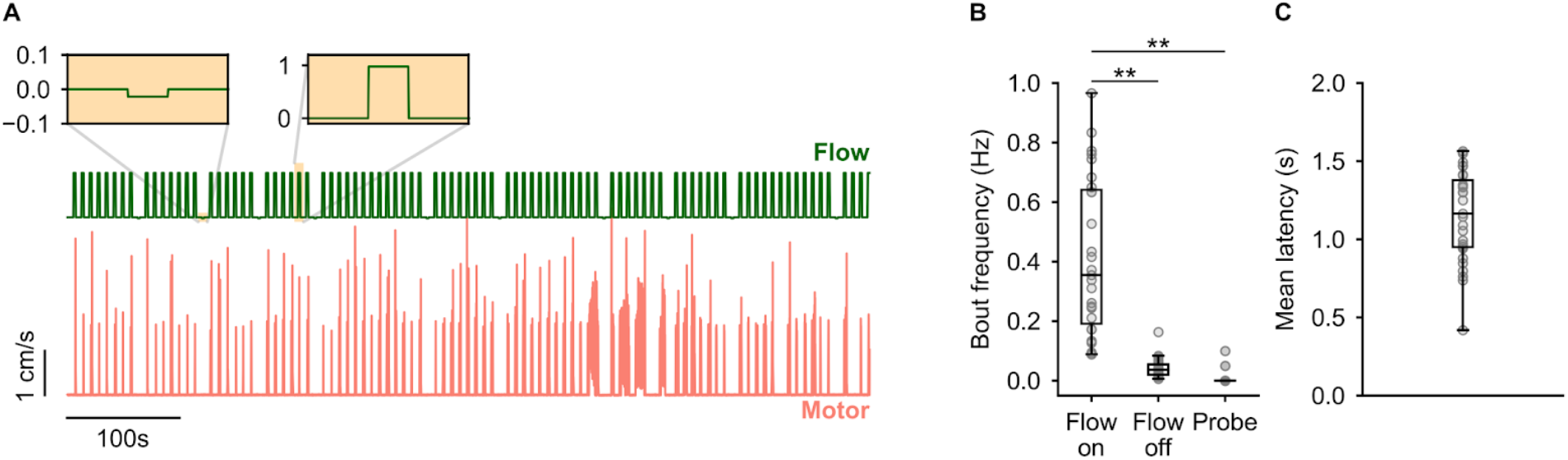
Experimental protocol to probe whether expectation of a stimulus influences behavioral responses. (A) Schematic of the 100 stimulus pulse long protocol. 10 negative probe stimuli were randomly interspersed amongst 90 swim-inducing 1cm/s forward optic flow stimuli. Zoomed insets show individual probe (left) and forward optic flow (right) stimuli. (B) Forward optic flow pulses of 2s duration reliably evoke swims. Almost no swims were seen in response to the probe stimulus or during the interval between stimuli. p = 1.01 x 10^-13^ by one-way repeated measures ANOVA, ** : p = 0.001 by Tukey’s post-hoc test, N=27 fish. (C) The mean latency with which swim bouts were initiated from the onset of forward optic flow stimuli.

